# Volumetric analysis of the terminal ductal lobular unit architecture and cell phenotypes in the human breast

**DOI:** 10.1101/2023.03.12.532249

**Authors:** Oona Paavolainen, Markus Peurla, Leena M. Koskinen, Jonna Pohjankukka, Kamyab Saberi, Ella Tammelin, Suvi-Riitta Sulander, Masi Valkonen, Larissa Mourao, Pia Boström, Nina Brück, Pekka Ruusuvuori, Colinda LGJ Scheele, Pauliina Hartiala, Emilia Peuhu

## Abstract

The major lactiferous ducts of the human breast branch out and ultimately end at terminal ductal lobular units (TDLUs). These glandular structures are the source of milk upon lactation, and also the most common origin of breast cancer. Despite their functional and clinical importance, the three dimensional (3D) architecture of TDLUs has remained undetermined, largely due to their absence in rodent animal models. By utilizing recent technological advances in optical tissue clearing, 3D reconstruction histology and microscopy, we imaged the 3D structure of healthy human breast tissue to determine whether general branching patterns or cell type specific niches exist in the TDLU. Our data demonstrate that highly branched TDLUs, also exhibiting elevated proliferation, are uncommon in the breast tissue regardless of donor age, parity or hormonal contraception. Overall, TDLUs have a consistent shape and their branch parameters are largely comparable between different TDLUs and individuals irrespective of donor age or parity. Simulation of TDLU branching morphogenesis in 3D by mathematical modelling suggests that evolutionarily conserved mechanisms regulate mammary gland branching in humans and mice, despite their anatomical differences. The data also demonstrate a new level of organization within the TDLU structure identifying a main subtree that dominates in bifurcation events and exhibits a more duct-like keratin expression pattern. In all, our data provide the first structural insights into 3D human breast anatomy and branching, and exemplify the power of volumetric imaging in deeper understanding of human breast biology.

**Graphical Abstract:** 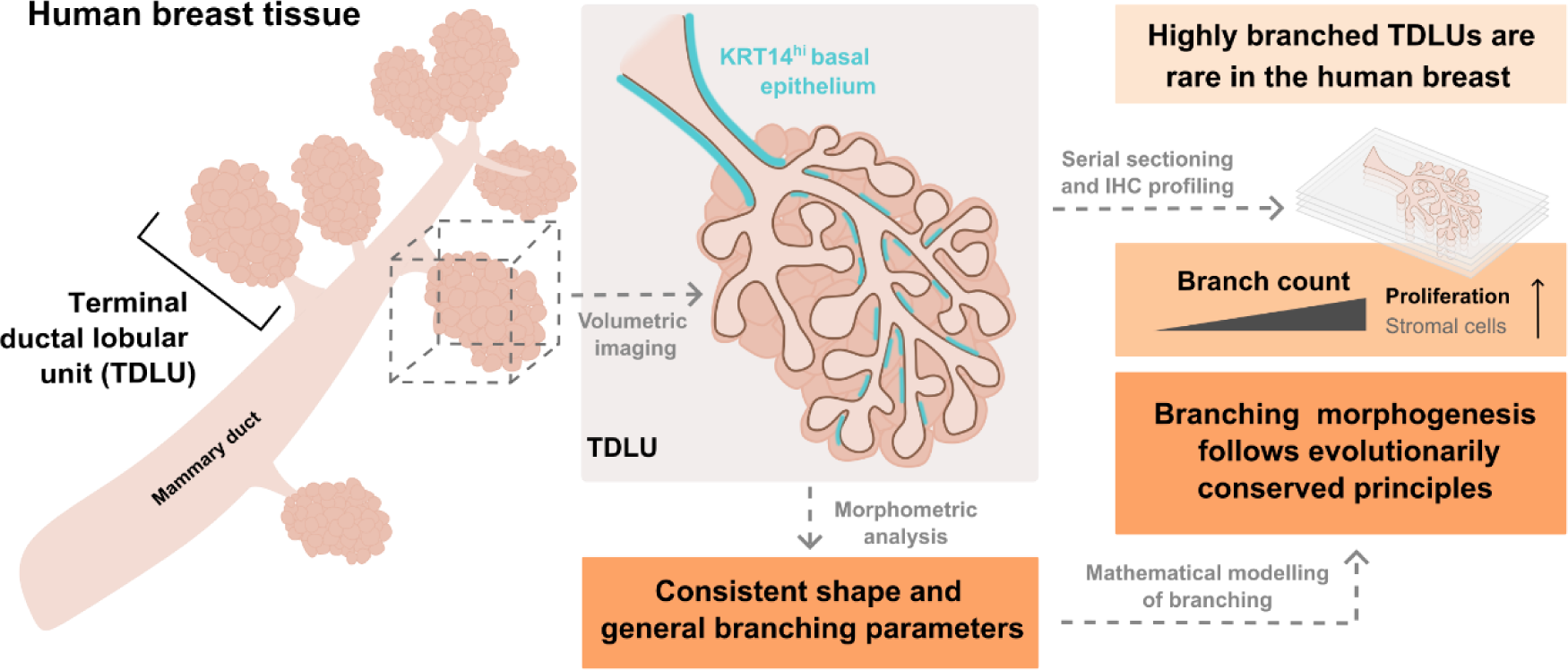

## Introduction

The human breast epithelium consists of a branched ductal network formed by 15-20 lobes in which collecting ducts, originating from the nipple, ultimately side-branch to form TDLUs (Bannister et al., 1995) (**Fig. 1A**). The majority of the gland forms during puberty, when in response to female systemic hormones such as growth hormone and oestrogen, the gland undergoes extensive branching morphogenesis laying down the main anatomical structures of the breast (Macias and Hinck, 2012). The morphology of the human breast is not static but changes dramatically during the reproductive cycles of pregnancy and lactation in order to adapt to the functional requirements of the tissue.

**Figure 1.**
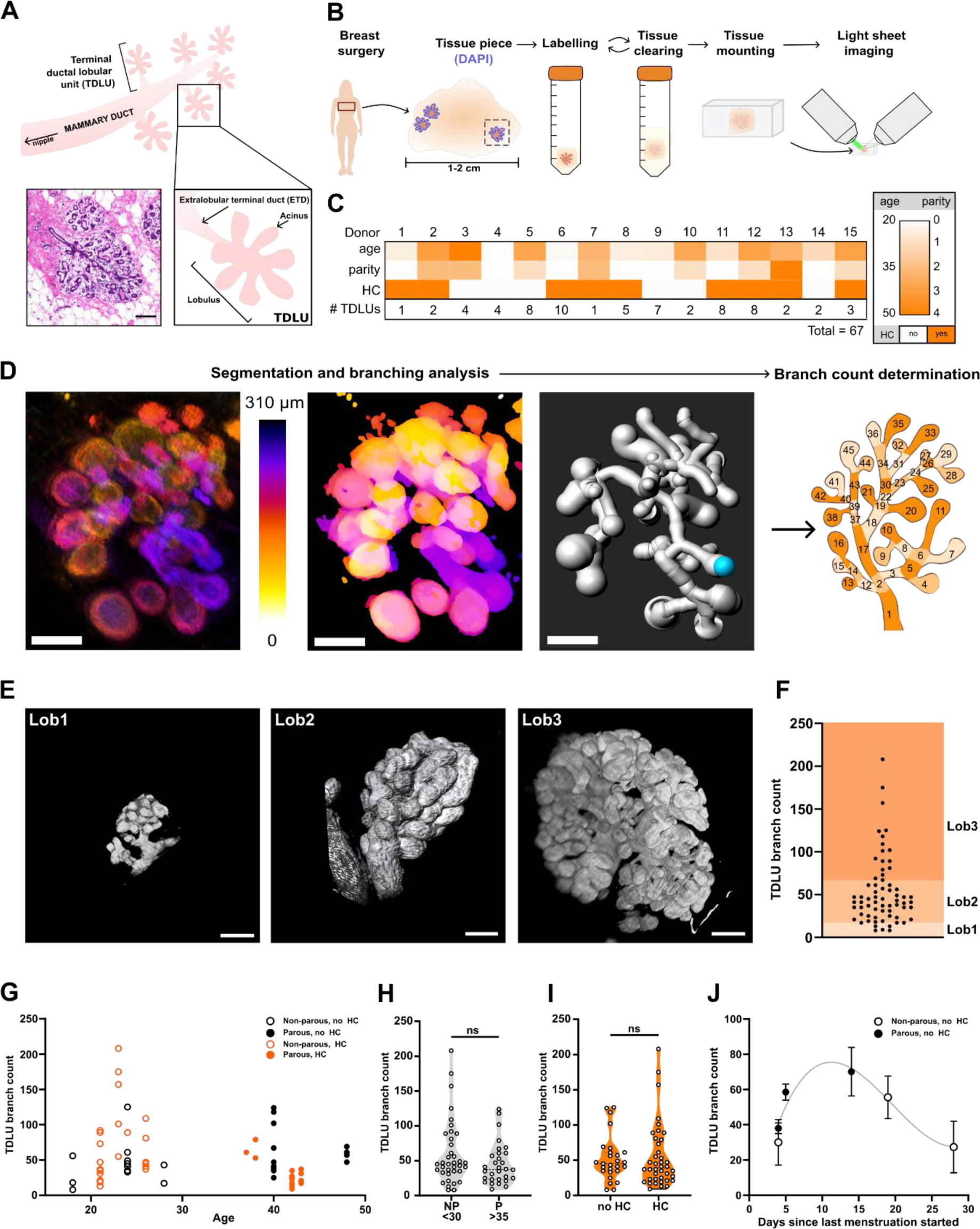
Volumetric imaging of optically cleared human breast tissue reveals that highly branched TDLUs are rare regardless of donor age and parity. **A.** A schematic representation of the human breast terminal ductal lobular unit (TDLU) based on current knowledge, and an H&E stained histological section of human breast TDLU (Donor 5). Scale bar: 100µm. **B.** Sample preparation and mounting protocol. Human breast tissue was dissected, fixed, and further trimmed to the smallest size possible utilizing DAPI signal at a fluorescence microscope. The pieces were immunolabelled and cleared for imaging. Finally, the tissue was mounted in agarose to be imaged with light sheet microscopy. **C.** The age, parity and use of hormonal contraceptives by the tissue donors. The number of TDLUs analysed per donor is indicated. **D.** Data analysis pipeline. A light sheet microscopy image of a TDLU visualized with depth colour coding (left), a segmented image visualized with the same color coding (middle), and an image of the branching structure model generated with Imaris FilamentTracer (right). Cyan dot in the image indicates the branch structure starting point. Scale bars: 100 µm. **E.** Representative images of breast tissue after iFLASH clearing and light sheet microscopy. Scale bars: 100 µm **F.** The number of branches per TDLU with respect to previous lobular typing (Russo et al., 1992). Lob1: 0-17 branches. Lob2; 35-60 branches. Lob 3; > 65 branches. **G.** The number of branches per TDLU graphed in relation to donor age. Parity of the donor or the use of hormonal contraceptives is indicated (n=67 TDLUs from 15 donors). **H-I.** The number of branches per TDLU graphed in relation to donor age and parity (**H**), hormonal contraceptive use (**I**) (n=67 TDLUs from 15 donors), or menstrual cycle phase (**J**) days since last menstruation started (n=26 TDLUs from 6 donors; mean ± SEM with non-linear curve fit).

TDLUs are considered the functional units of the breast and play a central role during milk production and lactation. Resting TDLUs are typically composed of an extralobular terminal duct (ETD) that branches into lobules with acini at the tips of the ducts (**Fig. 1A**). Interestingly, TDLUs are also considered the predominant anatomical location where breast cancer initiates (Wellings et al., 1975; Russo and Russo, 1997; Tabár et al., 2022). Therefore, detailed understanding of the structure and composition of the TDLUs is of high clinical importance, particularly related to their key role in breast cancer initiation.

While the use of rodent animal models has been pivotal for our basic understanding of healthy mammary gland development and structure, their usefulness for the study of TDLU formation is limited. Rodent mammary glands are formed by a single network of branched ducts within adipose stroma thus lacking the anatomically distinct lobes and TDLUs within a collagen-rich stroma (Dontu and Ince, 2015). Also, whether the branching mechanisms that produce the differential mammary gland outcomes are governed by the same principles in humans and mice has not been previously investigated.

Until now, human breast development, anatomy, and 3D structure have been described mostly based on low resolution imaging of histological dyes (Russo et al., 1994; Howard and Gusterson, 2000). While the histology and overall anatomy of the human breast epithelium, including the description of different TDLU categories based on donor age and parity, were defined (Russo et al., 1992), the 3D architecture and exact composition of TDLUs has remained obscure. Although much effort has been put into describing the epithelial and stromal cell populations in the human breast by single cell analysis (Nguyen et al., 2018; Pal et al., 2021; Chen et al., 2022; Gray et al., 2022), it is not known how these cell types are anatomically distributed within the breast epithelium and whether their abundance is linked to the extent of TDLU branching. Previous studies assessing cell localisation in breast tissue sections (Dontu and Ince, 2015; Villadsen et al., 2007; Kumar et al., 2023) or in micro-collected TDLUs/ducts (Goldhammer et al., 2022; Kohler et al., 2022) also provide limited information of the spatial distribution cell types within TDLUs. In addition, the extent to which TDLU structures vary between individuals and within a breast is unclear. As a result, TDLU composition and structure are still very ill defined.

As recent advances in technology now enable the imaging of larger intact tissue volumes labelled with multiple markers, we employed tissue clearing methods and volumetric light sheet microscopy to assess the 3D morphology of TDLUs derived from healthy breast tissue donors, both parous and nulliparous, in different age categories. We quantified TDLU structures and related their size and complexity to age, parity, and hormonal contraception status. Volumetric imaging was followed by multiplex immunofluorescence on the same TDLUs to relate cell type distribution and abundance to their 3D structure. Finally, we combined our quantitative analysis with mathematical modelling to investigate whether general branching patterns or principles exist for the human TDLU.

## Results

### Volumetric imaging of optically cleared human breast tissue reveals that highly branched TDLUs are rare regardless of donor age and parity

The morphology of entire human TDLUs has previously been documented only with bright field or confocal microscopy imaging thus lacking detailed volumetric information of the structure (Russo et al., 1992; Rios et al., 2019). In the early studies conducted on large numbers of breast tissue whole mounts using low resolution imaging, TDLUs were divided into four different types of branching, where lobular type 1 represents young nulliparous tissue (6-17 acini/TDLU), type 2 mature nulliparous tissue (17-60 acini/TDLU), type 3 parous tissue (>65 acini/TDLU), and type 4 intense branching occurring only during pregnancy and lactation (Russo et al., 1992). More recently, however, the number of acini per TDLU was reported to be comparable between nulliparous cases, and samples collected 18 months (Jindal et al., 2014), or 5 years (Ogony et al., 2022) postpartum.

To assess the influence of age, parity and hormonal contraception on the TDLU branch count using volumetric imaging, we obtained human breast tissue under informed consent from healthy premenopausal women of different age (18-48) and parity (0-4 children) undergoing breast reduction surgery (**Fig. 1B-C, Table S1**). Tissue pieces of approximately 5-20 mm in diameter were pre-screened for the presence of mammary epithelium by nuclear DAPI labelling, cleared with FunGI protocol (Rios et al., 2019) or with our improved version of the FLASH optical tissue clearing protocol (Messal et al., 2021) (termed hereon iFLASH), and labelled by immunofluorescence for cytokeratin 8 (KRT8) to visualize luminal mammary epithelial cells (**Fig. 1B**). We obtained good sample transparency and were able to visualize entire TDLU structures (**Fig. S1**). Altogether 67 TDLUs from 15 donors were imaged by volumetric light sheet imaging (**Fig. 1C**). From the obtained 3D image stacks, the TDLU structures were quantitatively analysed using Imaris FilamentTracer and TreeSurveyor softwares to visualise and quantify the branched networks (**Fig. 1D**). The luminal epithelium labelled with KRT8 formed a continuous tubular structure that bifurcated in seemingly random manner and ended as enlarged acini (**Fig. 1D-E**), demonstrating that the imaging protocol could reveal entire TDLU structures.

Quantification of the total 3D branch count for all analysed TDLUs indicated that the large majority of TDLUs were medium sized, corresponding roughly to type 2 lobules of earlier categorization (Russo et al., 1992)(**Fig. 1F**). The sample cohort was strongly divided into young (18-28) nulliparous and mature (35-48) parous (> 6 years postpartum or unknown) donors likely due to the relatively high average age of first-time mothers in Finland (30.1 years) (Perinatal statistics - parturients, delivers and newborns - THL, 2024) (**Fig. 1C, G, Table S1**). Importantly, the abundance of highly branched TDLUs was not higher in the samples from mature parous donors (**Fig. 1G-H**). In turn, the heterogeneity in TDLU branch count (**Fig. 1G**) was largest in samples from young nulliparous donors. More than half of the donors were using hormonal contraceptives but no emergent effect on TDLU branch count could be attributed to hormonal contraception (**Fig. 1I, Table S1**). Although based on a limited number of samples from women not using hormonal contraception, higher TDLU branch count was detected in the breast epithelium of women in the mid-phases of their menstrual cycle (**Fig. 1J**). Systemic estrogen levels, and thus mammary epithelial proliferation, have previously been shown to be higher approximately at this time of the cycle (Potten et al., 1988; Laidlaw et al., 1995). Overall, we generated a workflow for clearing and volumetric imaging of human breast tissue. Our data suggest that parity, in association with higher age, is not linked to larger number of TDLU branches (or acini) in the human breast tissue several years postpartum, thus confirming previous reports demonstrating similar findings based on tissue sections (Jindal et al., 2014; Ogony et al., 2022).

### TDLUs contain a main subtree that dominates in bifurcation events and exhibits a more duct-like KRT14 expression pattern

The location of different cell types in human breast tissue has been investigated using tissue sections (Dontu and Ince, 2015; Villadsen et al., 2007; Kumar et al., 2023) or in micro-collected TDLUs/ducts (Goldhammer et al., 2022; Kohler et al., 2022). However, the cellular composition or tissue-level organization in different parts of the TDLU remain poorly understood. KRT8 and cytokeratin 14 (KRT14) intermediate filament proteins have been established as luminal and predominantly basal epithelial markers, respectively, in the human mammary gland (Taylor-Papadimitriou et al., 1989; Gray et al., 2022). As expected, immunolabelling of these marker proteins in healthy breast tissue sections revealed higher KRT14 expression in the basal cell layer of the ETDs compared to most parts of the lobules (**Fig. 2A**).

**Figure 2.**
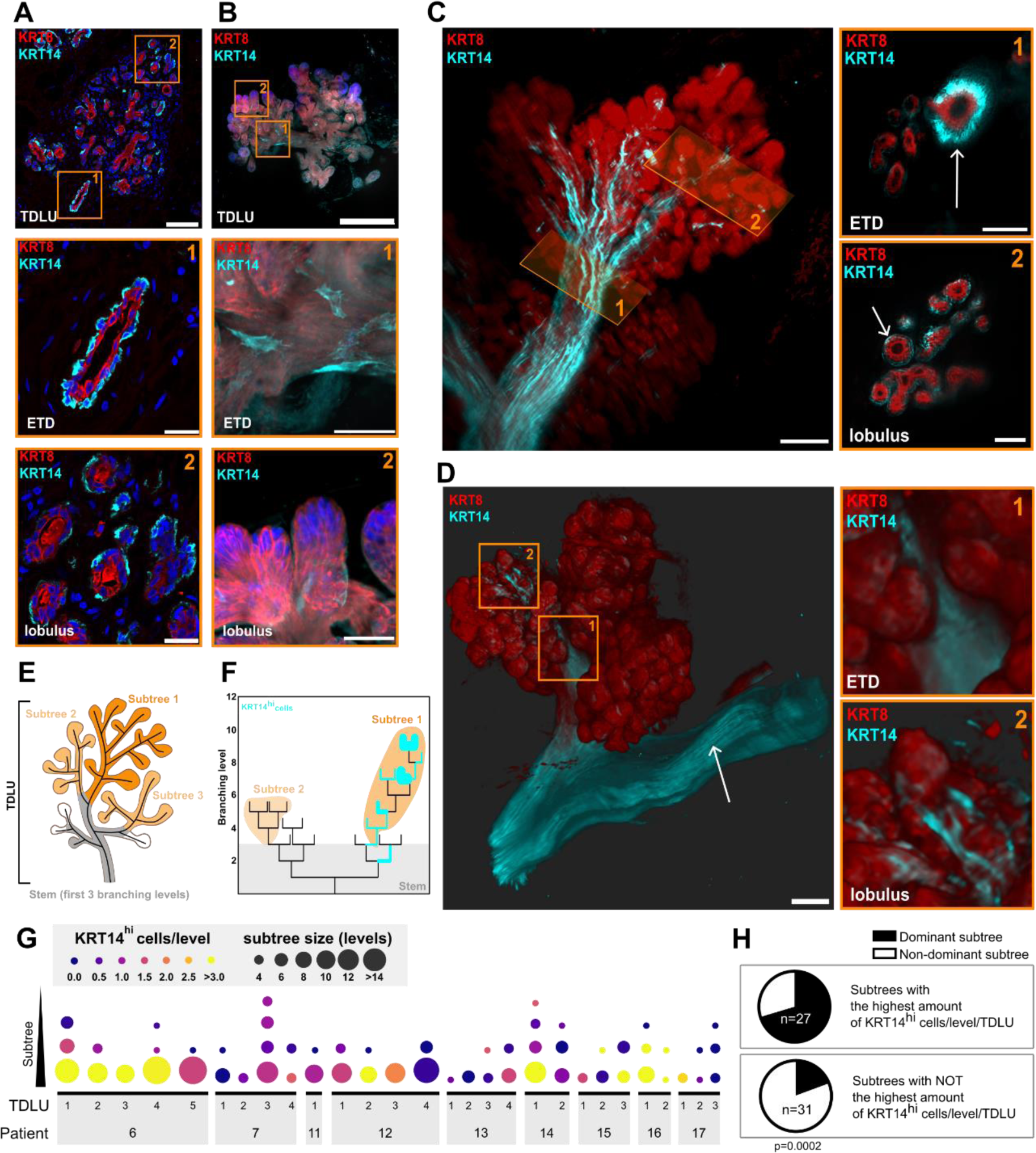
TDLUs contain a main subtree that dominates in bifurcation events and exhibits a more duct-like KRT14 expression pattern. **A-B.** Confocal microscopy imaging of KRT14 (basal epithelial, cyan), KRT8 (luminal epithelial, red) and nuclei (blue) in immunolabelled frozen human breast tissue sections (**A**) or in entire TDLUs of cleared breast tissue pieces (**B**). Higher levels of KRT14 are detected in the extralobular terminal duct (ETD) compared to the lobules of TDLUs. ROIs depict the ETDs (ROI 1) and lobuli (ROI 2). Scale bars: **A**: 100 µm (main), 25 µm (ROIs), **B**: 250 µm (main), 50 µm (ROIs). n=6-8 from 3 donors. **C-D.** Light sheet microscopy imaging of KRT14 (basal epithelial, cyan) and KRT8 (luminal epithelial, red) in entire TDLUs within cleared breast tissue pieces (**C**). KRT14^hi^ cells are visible in the duct (arrow), ETDs (ROI 1) and lobuli (ROI 2). Lower level of KRT14 is detected in the rest of the basal mammary epithelium (**D**). Scale bars: 100µm (main), 50µm (ROIs). Images are representative of 28 TDLUs from 9 donors. **E-F.** Determination of subtrees in TDLU branching structures (**E**). Representative tree structures of TDLUs with the distribution of KRT14^hi^ cells within the branching hierarchy (TDLU 5, cyan; line width is representative of the number of KRT14^hi^ cells in each branch. Branch length not to scale.) (**F**) **G**. Quantification of the branch levels and KRT14^hi^ cells in individual subtrees of the TDLUs demonstrating that the dominant subtrees retain high KRT14 levels (n=28, from 9 donors). Separate subtrees defined to begin at branch level = 3 and to persist for a minimum of 3 levels after cut-off). **H.** Quantification of the presence of KRT14^hi^ cells in main vs. not-main subtrees (n= 27-31 subtrees; Fisher’s exact test).

In order to build a more comprehensive view of the cellular organization of TDLU, we cleared breast tissue pieces with iFLASH protocol and immunolabelled the samples with KRT8 and KRT14 antibodies. The samples were imaged by confocal (**Fig. 2B**) or light sheet microscopy imaging (**Fig. 2C-D, Video S2**). Although some KRT14 expression was detected in the entire basal mammary epithelial cell layer (**Fig. 2C-D**), a subpopulation of basal cells expressing high levels of KRT14 (KRT14^hi^) was detected predominantly in the ETDs of whole TDLUs (**Fig. 2C-D**), as expected based on the tissue section analysis (**Fig. 2A**). Interestingly, volumetric imaging and quantitative analysis of the KRT14^hi^ basal cells present in the individual subtrees (**Fig. 2E-H**) revealed that the amount of KRT14^hi^ cells reduces gradually from the ETD towards the acini. Furthermore, the main subtree of a TDLU, defined as the subtree with most bifurcation events (levels), presents with a higher number of KRT14^hi^ basal cells per branch level (**Fig. 2G, Fig. S2**). In other words, the main subtree of each TDLU was significantly more likely to harbor the highest number KRT14^hi^ cells per branch level (**Fig. 2H**). Conversely, the expression of the luminal marker KRT8 seemingly increased in intensity towards the acini (**Fig. 2B-D**). Immunolabeling of another basal marker protein, transgelin, produced a clearly different expression pattern than KRT14, where expression levels were highest at the lobular level around the acini and lower close to the ETD (**Fig. S3A-B**). In all, these data reveal an additional level of organization within the TDLU structure identifying a main subtree that dominates in bifurcation events and exhibits a more duct-like KRT14 expression pattern, when compared to the smaller subtrees of the same TDLU.

### Multiplex immunohistochemistry of serially sectioned TDLUs demonstrates higher cell proliferation in highly branched TDLUs

In addition to the heterogeneity in protein expression levels, we also observed variation in TDLU size and branching complexity with branch counts ranging from 8 to 208 per TDLU (**Fig. 1F, Fig. S2**). The factors responsible for differential TDLU branching in the human breast remain unknown. To provide more quantitative insight into the tissue level features associated with different degrees of branching in human TDLU volumes (**Fig. 1**), we conducted serial sectioning of 110 TDLUs from samples previously imaged by light sheet microscopy (**Fig. 1-2**), and immunolabelled the samples with multiplex panels of antibodies for known mammary gland cell markers (**Fig. 3A**). H&E staining was performed at a 40 µm interval (every 10^th^ tissue section) for the entire tissue sample volumes (**Fig. 3B**) which could also be processed for 3D reconstruction histology to visualize the structure in 3D (**Fig. 3C**). Calculation of the cumulative branch count from the H&E sections for each TDLU resulted in a comparable distribution of the branch counts per TDLU (**Fig. 3D, Fig. S4**) as was detected by light sheet imaging (**Fig. 1F**), even though the number of TDLUs that could be visualized from the serial sections was higher. The categories from low to high TDLU branching were based on the quartiles of branch count per TDLU in the whole data set (1^st^ - 4^th^ quartile equivalent to ranges of 0-25%, 25-50% 50-75% or 75-100% in branch count) (**Fig. 3D**).

**Figure 3.**
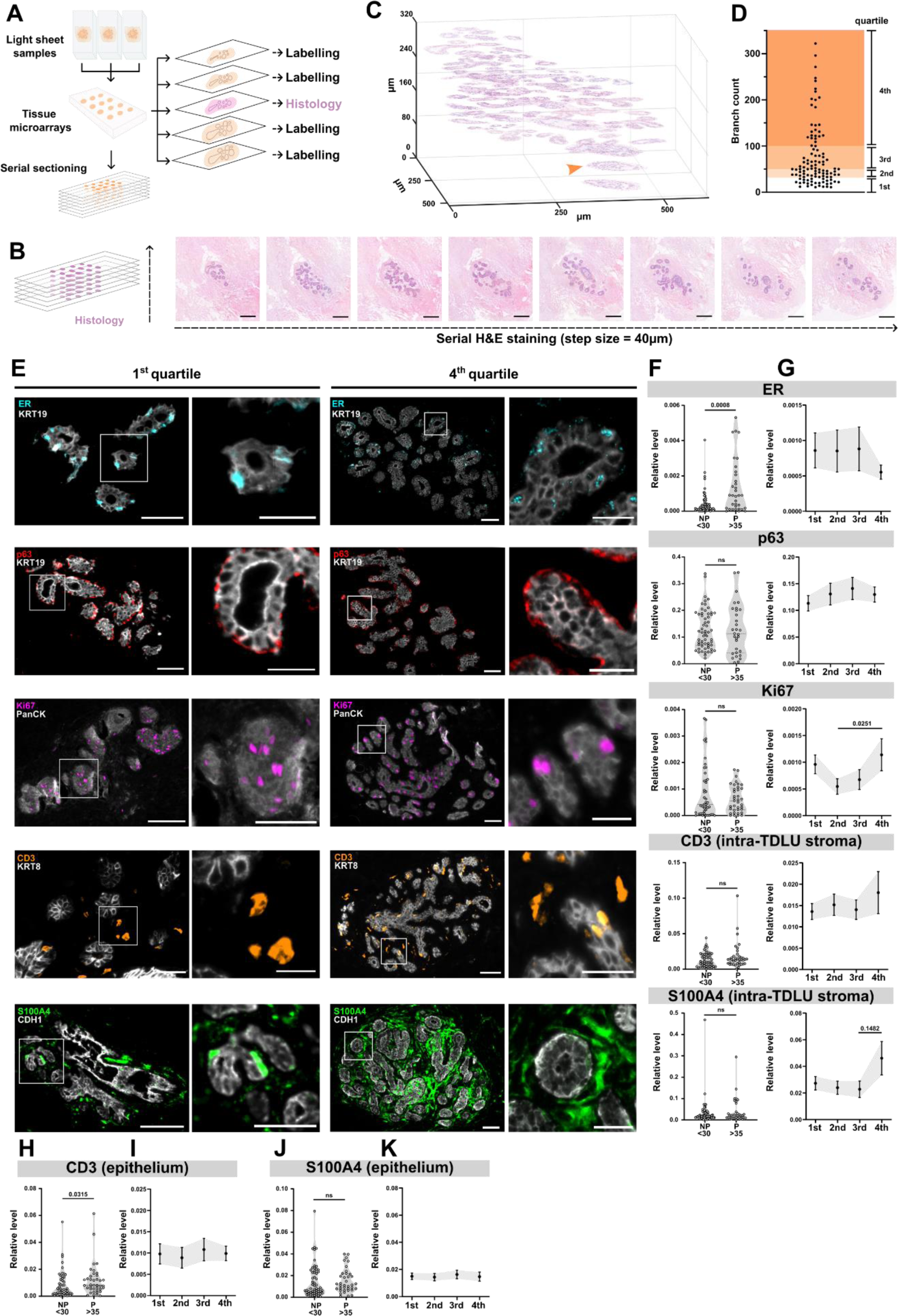
Multiplex immunohistochemistry of serially sectioned TDLUs demonstrates higher cell proliferation in highly branched TDLUs. **A.** Serial section sample preparation protocol. Tissue samples utilized for volumetric light sheet imaging were re-embedded into tissue microarrays (TMAs) and serially sectioned (4µm sections). Every 10^th^ section was H&E stained, allowing for 3D reconstruction of tissue anatomy. Remaining, consecutive sections were used for multiplex labelling. **B-D.** An example of H&E series of a TDLU (**B**). 3D reconstruction histology of the structure in (**C**). ETD (orange arrow). **D.** Distribution of all analyzed TDLUs based on branch count calculated from the H&E stained serial sections (n=110 from 10 donors). Scale bars (B): 200µm. **E.** OPAL 7-plex immunolabelling with antibodies against ER (cyan), p63 (red), Ki67 (magenta), CD3 (orange) and S100A4 (green). Scale bars: 50 µm (main), 25 µm (ROIs). **F-K.** Quantification of specific IHC signals/cell types per TDLU in function of donor age/parity (**F, H, J**) or in function of the percentiles of branch count (**G, I, K**). Level of signal of interest was related to applicable tissue area; ER to luminal epithelium (KRT19), p63 to basal epithelium (αSMA), Ki67 to epithelium (pan-keratin), CD3 to within [epithelium] or outside of luminal epithelium [intra-TDLU stroma] (KRT8) and S100A4 to within [epithelium] or outside of epithelium [intra-TDLU stroma] (CDH1).

The OPAL 7-plex immunolabelling panels were labelled at 2-3 z-positions per sample array and contained a range of established markers for the mammary epithelium [E-cadherin (CDH1), pan-keratin], luminal epithelial cells [KRT8, KRT19, estrogen receptor (ER)], basal myoepithelial cells [p63, α-smooth muscle actin (αSMA, not shown)] (Prabhu et al.; Kumar et al., 2023), T-cells (CD3), stromal cells (S100A4, also known as FSP-1) (Fendt et al., 2023) and cell proliferation (Ki67) (van Dierendonck et al., 1989) (**Fig. 3E**). The samples were visualised by widefield fluorescence microscopy and individual

TDLUs analysed quantitatively for the relative amount of selected markers in the epithelial or stromal areas. The level of each marker in the TDLUs was correlated with the parity and age of the donor (**Fig. 3F**) and the quartiles of branch count (**Fig. 3G**).

ER was expressed in a subset of KRT19^+^ luminal cells, as expected, and exhibited no clear correlation with the TDLU branch count, although a trend towards lower number of ER^+^ luminal cells was observed in highly branched TDLUs (**Fig. 3E, G**). In turn, the amount of ER^+^ cells was higher in the mammary epithelium of mature parous donors compared to young nulliparous (**Fig. 3F**), as was previously shown to occur during aging (Shoker et al., 1999). On the contrary, p63^+^ basal myoepithelial cells were detected in equal amounts in the human TDLUs irrespective of age, parity, or TDLU branch count (**Fig. 3E-G**). Interestingly, proliferation was higher in TDLUs with highest branch count (4^th^ quartile) compared to TDLUs with lower branch count (2^nd^ quartile) as demonstrated by immunolabelling of Ki67 (**Fig. 3E-G**), suggesting that highly branched TDLUs could be in a more proliferative state compared to an average-sized TDLU.

In the stromal compartment of the TDLU, we observed stromal CD3^+^ T-cells in equal amounts irrespective of age, parity, or TDLU branch count (**Fig. 3E-G**). Interestingly, mature and parous women had more intraepithelial CD3^+^ T-cells than young nulliparous donors (**Fig. 3H**), but the amount of intraepithelial CD3^+^ T-cells was not correlated to TDLU branch count (**Fig. 3I**). An interesting trend, although not statistically significant, was observed in stromal S100A4^+^ cells, which marks fibroblasts, T cells, monocytes, and macrophages (Fendt et al., 2023). S100A4 labelling was somewhat higher in the stroma of TDLUs with highest branch count (4^th^ quartile) compared to TDLUs with lower branch count (**Fig. 3E-G**). Low amount of S100A4^+^ cells was detected among the epithelium regardless of TDLU branch count (**Fig. 3J-K**). The finding that highly branched TDLUs appear more populated by S100A4^+^ cells is well in line with previous studies demonstrating that the presence of fibroblasts and macrophages promotes the branching morphogenesis of mammary epithelial cells *in vitro* (Sumbal and Koledova, 2019; Sumbal et al., 2024), and *in vivo* in mice (Gouon-Evans et al., 2000). The exact nature of the S100A4^+^ cells remains to be investigated further. Together, our data provide the first evidence for clinical relevance of the regulatory role of stromal cells in the branching morphogenesis of the human TDLU.

### Volumetric imaging of optically cleared human breast tissue reveals consistent branching parameters between different TDLUs and individual donors

The relatively high variation in TDLU branch count prompted us to investigate the morphological similarity of individual branches in TDLUs. In order to quantitatively establish the 3D morphology and dimensions of the human TDLU, we wanted to avoid distortion by tissue shrinkage, and used optical tissue clearing with FUnGI protocol that, unlike several other protocols, has been reported to preserve the original size of the specimen (Almagro et al., 2021; Rios et al., 2019). Breast tissue samples from four donors were cleared with FUnGI, and 14 TDLU structures were successfully imaged with volumetric light sheet microscopy (**Fig. 4A**). The donors represented different ages (range 24-50 years) and parity (range 0-2 children) (**Table S1, Fig S5**). For quantitative branching analysis, the 3D image stacks were segmented for KRT8 signal using WEKA trainable segmentation in ImageJ followed by quantitative analysis of the TDLU structures using Imaris FilamentTracer (**Fig. 4B-C, Video S1**).

**Figure 4.**
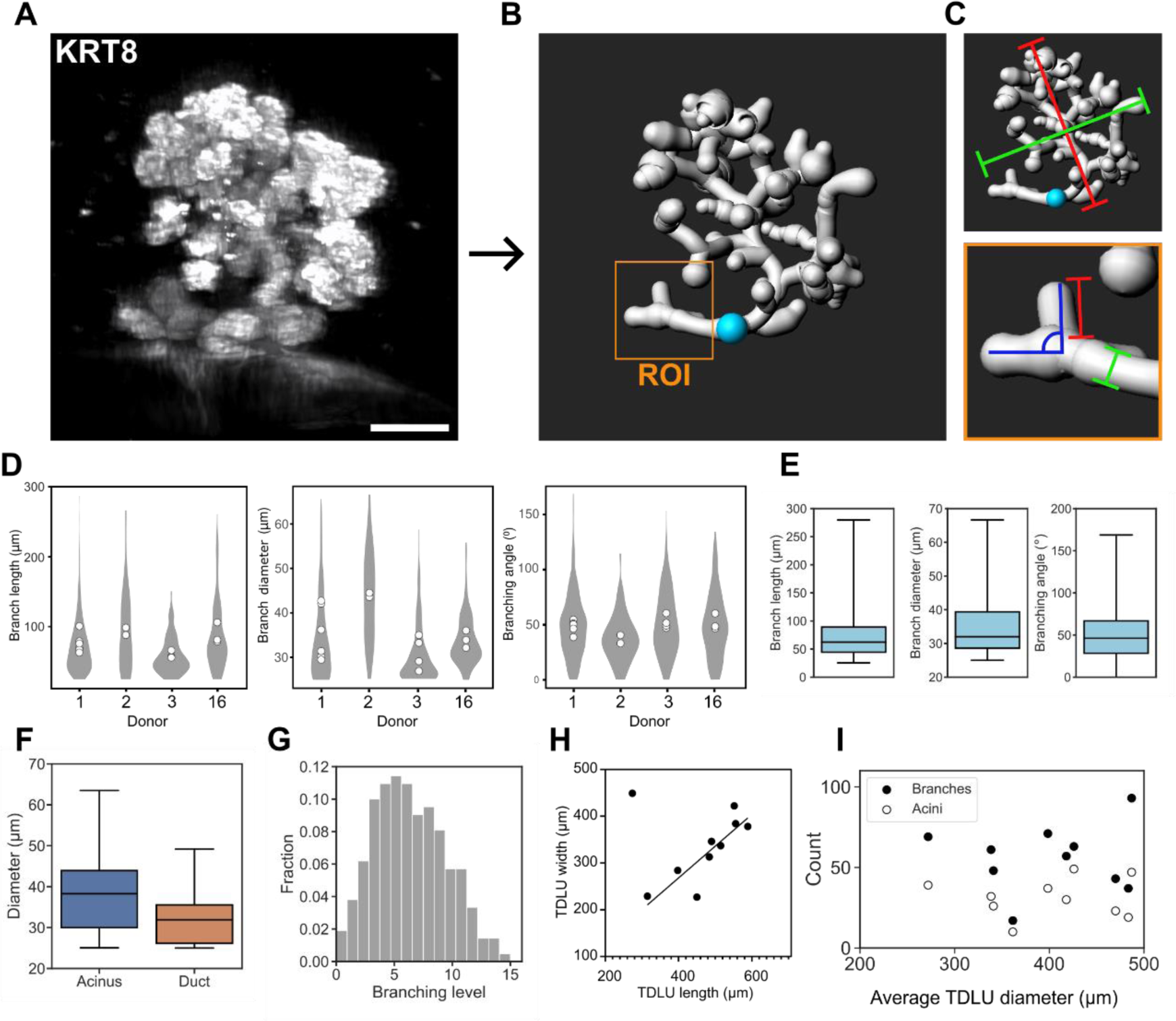
Volumetric imaging of optically cleared human breast tissue reveals consistent branching parameters between different TDLUs and individual donors. **A-B.** Representative image of breast tissue cleared with FunGI, imaged with light sheet microscopy (**A**) and with the branch architecture segmented for quantitative branch analysis (**B**). Scale bar: 100µm. **C.** Parameter definitions. Top image: TDLU length (red line) and TDLU width (green line), cyan dot indicates the network starting point (ETD).Bottom image: branch length (red line), branch diameter (green line), and branching angle (blue). **D.** Distributions of the parameters branch length, branch diameter and branching angle by donor. Mean values of individual TDLUs are shown by the white dots and grey area shows value distribution. **E.** Pooled distributions of branch length, diameter, and angle (n_length_=689, n_diameter_=689, n_angle_=689 branches from 14 TDLUs of 4 donors). Boxes show median and interquartile range, and whiskers the full range of data points. **F.** Distributions of acinar and ductal diameters per TDLU (n=14 from 4 donors). **G.** Distribution of branching levels. Data is pooled from all complete TDLU structures for which the whole network could be reliably traced (n=8 TDLUs from 4 donors). **H.** Measured length and diameter of the whole TDLU. Line shows linear fit to data excluding the outlier point. TDLU aspect ratio from the fit is 1.47 (n=10 TDLUs from 4 donors). **I.** Total number of branches and tip branches (acini) in TDLU as a function of average TDLU diameter. Average TDLU diameter is the average of the measured TDLU length and width. (n=10 TDLUs from 4 donors).

The results of the morphometric analysis of human breast TDLUs demonstrate regularity in branch (also referred to as ductules) length, diameter and angle (**Fig. 4D-E, Fig. S6B-D**). The average branch length was 71 µm (s.d. 37.8 µm), diameter 35 µm (s.d. 8.9 µm), and the average branch angle with respect to the parent duct was 49° (s.d. 29.9°) (**Fig. 4E**). The TDLU tip branches (acini) were on average 1.2 times as wide in diameter as the ducts in the TDLU (**Fig. 4F**). A TDLU formed most commonly from 4-6 consecutive bifurcation events, and branches at levels higher than 10 consecutive bifurcations were few (**Fig. 4G**). Average TDLU length (distance from ETD to the farthest acinus, **Fig. 4H**) and width (largest width in the direction perpendicular to the ETD direction, **Fig. 4H**) were 460 µm and 360 µm, respectively, and TDLUs typically have a regular shape with an aspect ratio of 1.5 (**Fig. 4H**). These measures are similar to the previously reported size of a normal TDLU based on HE stained tissue sections (Figueroa et al., 2014). Interestingly, the branching of normal mammary epithelium adjacent to a premalignant lesion (ductal carcinoma *in situ*) was comparable to healthy donor tissues (**Fig. 4D**, Donor 16). Our data also demonstrate that TDLUs have a consistent shape and their branch parameters are largely comparable between different TDLUs and individual donors of variable age and parity (**Fig. 4D**). In contrast to previous observations using tissue sections (Rosebrock et al., 2013), the total number of branches or acini (tips) per TDLU was not proportional to the TDLU diameter (**Fig. 4I**), suggesting that branching density may vary.

### Human mammary gland branching morphogenesis appears to be governed by the same principles as in mouse

Our volumetric analysis highlighted that despite several common features, such as the existence of a dominating TDLU subtree (**Fig. 2**), and the consistent TDLU shape and branch parameters (**Fig. 4**), human mammary gland TDLU branching is irregular, asymmetric, and variable in branch count (**Fig. S7, Fig. S6E**). A fractal branching model was previously proposed for the morphogenesis of human mammary gland TDLUs (Honeth et al., 2015), but the symmetrically occurring bifurcations and the gradually decreasing branch length and diameter required by this model do not capture the 3D branching structure of the human TDLU (**Fig. S6B-D**). The fact that human TDLU branching networks are not self-similar, ie. each structure is unique and varies greatly in size (**Fig. S4, Fig. S6E, Fig. S7**) also speaks against a stereotyped branching morphogenesis observed for example in the lung (Metzger et al., 2008) or in the kidney (Lefevre et al., 2017), and the absence of loop structures does not support models where the final tree-like form is reached through remodelling of a mesh of interconnected ducts (Dahl-Jensen et al., 2018).

Branching morphogenesis by branching and annihilating random walk (BARW) has previously been used to simulate generation of mouse mammary gland (Hannezo et al., 2017; Nerger et al., 2021), kidney (Hannezo et al., 2017), pancreas (Sznurkowska et al., 2018), zebrafish nervous system (Uçar et al., 2021) and cholangiocyte organoids (Roos et al., 2022). Whether human TDLUs branch and develop with the same principles as the mouse mammary gland has not been addressed. In the mouse mammary gland BARW model, branches grow forward, bifurcate randomly with a constant probability, and terminate if they grow too close to an existing branch (Hannezo et al., 2017). Growing branches avoid collisions to other branches which is modelled as repulsion from branches (Uçar et al., 2021).

Given that the mouse mammary fat pad is very flat, the simulation area was defined in 2D (Hannezo et al., 2017). In order to adapt the BARW model to human TDLUs that are smaller and 3D compared to murine mammary gland, we defined that bifurcation happens to a random 3D orientation angle in 0-360° (**Fig. S6F**) and with an opening angle drawn from our experimentally observed distribution (**Fig. S6G**). For the mathematical modelling of human TDLU branching, we also utilized the experimentally obtained parameters for branch diameter, bifurcation probability, termination distance, repulsion distance, and branching and rotation angle distributions,(**Fig. 4E, Fig. S6F-G**, **Table S2**) as explained in Materials and Methods. Value of the branch repulsion strength was optimized by finding the best match between the branch count distributions of the simulated and experimentally observed networks. A branching free zone of radius equal to branch diameter was defined around each branching event to prevent immediate re-branching which sets a requirement that branches shorter than their width are not mature enough to branch again (**Fig. 5A**) (Roos et al., 2022).

**Figure 5.**
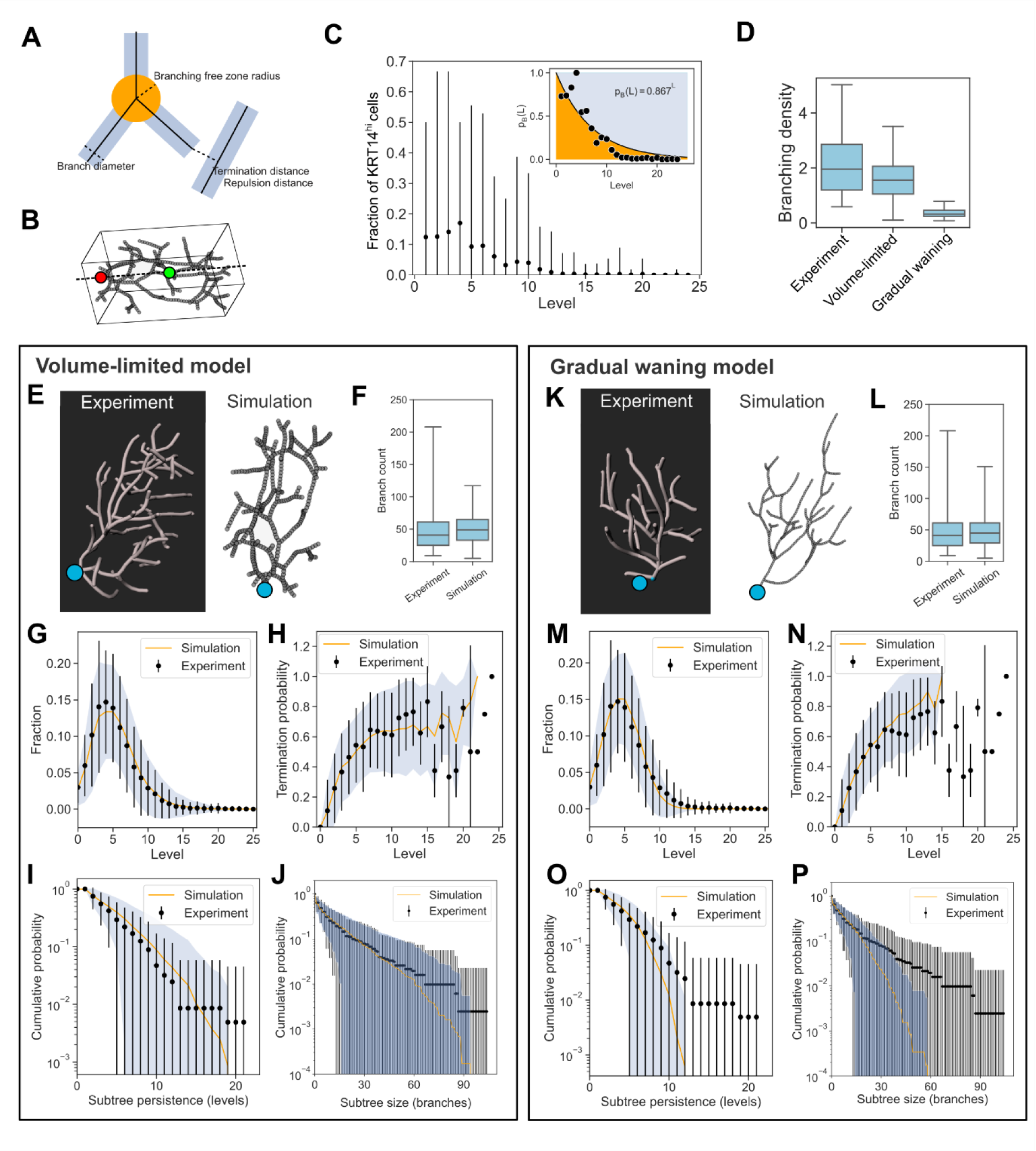
Simulation of human TDLU branching morphogenesis with volume-limited BARW model resembles more accurately the experimental data. **A.** Schematic image drawn to scale visualizing simulation parameters. **B.** Simulation volume control in volume-limited model. Red dot marks the network origin and green dot the center of mass of all network points. Dashed line indicates the general direction of the network, defined by a vector from network origin to center of mass. **C.** Simulation volume control in gradual waning model. Main image shows the fraction of KRT14^hi^ cells of all the KRT14^hi^ cells in the TDLU by levels. Dots mark the average value and error bars the variation of all TDLUs. Inset shows the fit of the exponential equation to the experimental KRT14 cell distribution used in the model as the waning boundary between branching (orange area) and self-termination (blue area). n=28 TDLUs of 9 donors. **D.** Branching density for experimental TDLUs and both simulation models. Branching density is calculated as the number of branches per whole network volume, and it is given in units 1/(10^6^ µm^3^). n_Experiment_=8 TDLUs of 4 donors, n_Simulation_=1000 for each model. **E, K.** Images of experimental and simulated networks for both models. Blue dot indicates the network beginning point. Line and point sizes are not to scale. **F, L.** Simulation predictions for the number of branches in a TDLU compared to experimental results. **G, M.** Distribution of branching levels in experimental data and simulation results. **H, N.** Probability of branch termination at different branching levels. **I, O.** Probability of a subtree beginning at level 3 to span a number of levels. **J, P.** Probability of a subtree beginning at level 3 to contain a number of branches. In G, H, I, J, M, N, O, and P dots and error bars indicate mean and standard deviation of the experimental data, and the orange line and blue shaded area mean and standard deviation of results of 1000 simulations. for **F-J** and **L-P** n_Experiment_=63 TDLUs of 15 donors, n_Simulation_=1000 for each model.

In the simulations growth takes place mostly in the perimeter of the network by branches invading into empty space, and the networks will grow indefinitely unless growth is somehow limited. We tested two mechanisms for limiting the network growth: 1) Growth was limited to a predefined volume (Hannezo et al., 2017), and 2) a model in which branches have gradually waning growth potential that eventually stops the branching. The latter model was inspired by the accumulation of KRT14^hi^ cells into one TDLU subtree that reached higher branch levels (**Fig. 2**), which could indicate the existence of unequal and potentially reducing growth capacity. In the volume-limited model we let the network grow within a volume equal to an experimentally observed average TDLU volume of 460 µm × 310 µm × 310 µm. Any branch reaching the edges of the simulation volume was terminated. Instead of rigid maximum limits for *x*, *y*, and *z*, we used an adaptive limiting volume (**Fig. 5B**) that follows the network growth direction to maximize the use of available space by preventing occasional premature termination by all branches colliding directly to network volume edges, which may happen if the whole network starts to grow towards one side of the volume.

In the model of gradually waning branching potential, we utilized a rule that instead of always branching at the end of the branch growth phase (as in the original BARW model), the branch has a branching level dependent probability to self-terminate. Self-termination represents a situation where the branch seizes to grow, no more branching is instigated and the branch becomes inactive. At level 0 the probability to branch is 1 and the balance shifts towards self-termination in higher branching levels (**Fig. 5C**). Exponential equation *p*_*B*_(*L*) = *q*^*L*^ was used to represent the probability to branch describing fractional decline in branching potential with branch level *L*. Probability to terminate was *p*_*T*_ = 1 − *p*_*B*_. Hypothesizing that KRT14^hi^ cells could represent a feature of more duct-like cells with higher branching potential, we estimated the decline rate *q* = 0.867 from the experimentally observed decline of the KRT14^hi^ cell distribution in the TDLUs (**Fig. 5C**).

Simulation with the volume-limited model produced ductal structures very similar to the TDLUs in human tissue samples (**Fig. 5E-J** and **Fig. S7**). Simulated networks had similar branch counts to those observed experimentally (**Fig. 5F**), and the networks were equally effectively filling the space available to them (**Fig. 5D**). Model prediction of branch level distribution followed the general trend of that observed experimentally (*R^2^* = 0.99, **Fig. 5G**). Ratio of number of terminal branches (acini) to total number of branches in simulated branching networks is 0.51 (**Fig. S6A**) matching well the experimentally observed value 0.55. This is expected since in a bifurcating network at any given time point the number of active tips at branching level *L* is 2^*L*^, and the total number of branches follows a geometric series 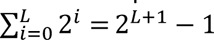. The ratio of the number of acini to the total number of branches is then 2^*L*^⁄(2^*L*+1^ − 1), which is 0.5 at large *L*. Ratios slightly higher than 0.5 seen in experimental and simulation data are explained by the terminating branches and side branching.

Topology, ancestry of the subtrees, and connectedness of the networks was compared by calculating the branch termination probability, and sizes of subtrees in levels and branch counts (**Fig 5H-J**) as before (Hannezo et al., 2017; Roos et al., 2022). Probability of a branch terminating at a given level (**Fig. 5H**) follows a similar trend as observed before (Hannezo et al., 2017; Roos et al., 2022) levelling to approximately 0.6 at levels above 5. Levelling happens at earlier levels compared to e.g. mouse mammary gland due to the lower branch count in human TDLUs, which also explains the rising probability at higher levels. Simulated networks follow the trend well (*R^2^* = 0.90, **Fig. 5H**).

Tree plots of the branch ancestry (**Fig. S7**) show that human TDLU subtrees are not self-similar or symmetric, but instead their size varies substantially within a TDLU. Many shallow subtrees are typically observed at low branch levels (spanning only 1-5 levels from the root point) and highest branching occurs in 1-2 ancestral subtree lines that continue growing up to levels 10 and above (**Fig. S7**). Comparable behaviour can also be generated by the simulations (**Fig. S7**) which is quantitatively shown using analysis of subtree sizes (**Fig. 5I-J**) (Hannezo et al., 2017). The fit of model results to experimental observations is good for subtree persistence (*R^2^* = 0.99, **Fig. 5I**), and for subtree size in branches (*R^2^* = 0.99, **Fig. 5J**). Overall the 3D simulation based on the BARW model with limiting volume produces a close fit with the experimental data.

The model of gradually waning branching potential was less successful in reproducing the features of the experimental networks. If the networks were allowed to grow freely, their shape was more spherical or even flatter with aspect ratio < 1 (**Fig. 5K**). To reproduce the realistic network shape with aspect ratio of 1.5, another model parameter, a global guidance force, had to be introduced (Uçar et al., 2021). With an optimized value of 0.1 for the global guidance strength parameter the realistic shape could be reproduced without affecting the network topology results. In general, this model produced also a satisfactory fit to the experimental data (**Fig. 5L-P**). For example, branch counts were equal (**Fig. 5L**). However, these networks growing in empty space grew much larger in volume, as indicated by the much less densely packed structure (**Fig. 5D**). The model was neither able to produce networks in which subtrees would persist to highest levels above 15 (**Fig. 5O**) and subtrees also consisted of fewer branches (**Fig. 5P**) even if the overall branch count distribution was similar to that observed experimentally (**Fig. 5L**). This indicates that growth in these networks takes place in several shallow subtrees growing in parallel, rather than 1-2 main subtrees that have most of the branches and extend to high branching levels as was observed experimentally, and better modelled by the volume limiting model (**Fig. S7**).

In summary, the simulations demonstrate that branching morphogenesis of human TDLUs appears to be governed by the same principles as in mouse, with the BARW model adapted to a 3D volume capturing well the morphology of the experimentally analysed TDLU structures.

## Discussion

The 3D branching structure of the human breast has remained largely unexplored, particularly in high resolution and in a quantitative manner. Here, we set out to provide the first structural insights into the 3D anatomy, cell type location and branching of resting human TDLUs. While TDLUs of different donors were heterogeneous in branch count, they had a consistent shape and branching parameters, such as branch length, width and angle. Our data provide evidence that resting TDLUs of young nulliparous women are indistinguishable from TDLUs of mature parous donors in terms of branch count, but have lower amounts of ER-positive luminal epithelial cells and intraepithelial T-cells. While hormonal contraception did not seem to influence the overall branch count of TDLUs in our sample cohort, branching varied during the menstrual cycle, as expected. Highly branched TDLUs had more proliferating epithelial cells compared to less branched and, interestingly, exhibited a trend towards higher amount of stromal cells surrounding the epithelium. Our data also demonstrate a new level of organization within the TDLU structure identifying a main subtree that dominates in bifurcation events and exhibits a more duct-like KRT14 expression pattern. Finally, our data suggest that human TDLUs form by evolutionarily conserved mechanisms that regulate mammary gland branching both in humans and mice.

We simulated branching morphogenesis of the human TDLU by adapting the BARW model (Hannezo et al., 2017) to 3D, and using experimentally defined parameters for the modelling. Two potential mechanisms were considered for limiting the network growth: 1) limitation of growth to a predefined volume, as previously used in the modelling of mouse mammary gland branching (Hannezo et al., 2017), or 2) limitation of growth by gradually waning growth potential. The latter mechanism was based on the gradual decline we observed in KRT14^hi^ cells towards the acini of the TDLU. KRT14 has previously been linked to increased adhesive and migratory properties (Fujiwara et al., 2020; Cheung et al., 2013), as well as mechanical integrity of cells (Ramms et al., 2013). Thus, we hypothesized that KRT14 expression level could potentially reflect local differences in growth or migration potential, and presumed that a decline in branching potential could provide sufficient mechanism to regulate TDLU growth. While the simulated networks resulting from mathematical modelling with this mechanism for limiting the network growth had similarities with the experimental data, they were sparse and lacked a dominating subtree.

Instead, the networks simulated with the volume-limited BARW model resembled highly the TDLUs in the clinical specimen despite the fairly wide variation in the data, suggesting that human mammary gland branching morphogenesis is governed by the same principles as the mouse mammary gland. The most significant difference between the species in terms of the modelling parameters is the branching area which in humans was limited to a single TDLU. Our data point to the fact that the TDLU form is likely limited by interaction with the stroma. Why TDLUs evolved and why they adopt this particular size and shape in humans remains unknown. Furthermore, whether the morphogenesis of an entire lobe would follow comparable principles with TDLU branching remains to be studied. Specifically, it would be interesting to address, if the larger branch diameter in collecting ducts is sufficient to account for the higher branch length.

Terminated mammary ductal tips typically reside close to an existing duct or the fat pad boundary in mice (Silberstein, 2001). Additionally, ductal branching was shown to occur *in vitro* only when the tip was remote from the other ducts (Nelson et al., 2006). Consequently, ductal elongation and branching was suggested to proceed as a ‘‘default state’’ with tip termination occurring only when tips come into proximity with existing ducts or the edges of the fat pad (Hannezo et al., 2017). This scenario, as well as the modelling presented here, suggest that growth limiting mechanisms exist around the TDLUs as well. While there is currently limited information on the negative regulation of branching morphogenesis, TGFβ1 secretion (Silberstein and Daniel, 1987; Nelson et al., 2006) and progressive envelopment of the gland by extracellular matrix (Silberstein and Daniel, 1984; Osin et al., 1998) have been suggested to limit mammary gland branching. Therefore, gradual alterations in TDLU proximal stroma may contribute to TDLU growth inhibition.

Mammary gland stromal cells have been shown to promote mammary epithelial branching (Gouon-Evans et al., 2000; Sumbal and Koledova, 2019; Sumbal et al., 2024). We observed a trend towards an elevated amount of S100A4^+^ stromal cells in the TDLUs with highest branch count where also proliferation was higher than in less branched TDLUs. S100A4 (also known as FSP-1) expression has been detected for instance in fibroblasts (Cheng et al., 2005; Trimboli et al., 2009; Pickup et al., 2015; Koledova et al., 2016) and macrophages (Le Hir et al., 2005; Österreicher et al., 2011). Two distinct fibroblast lineages, lobular and interlobular, have been described in the human breast (Morsing et al., 2016). Intralobular fibroblasts, also quantified in our study, were shown to support epithelial growth and morphogenesis (Morsing et al., 2016). FGF signaling has been demonstrated to regulate the ability of stromal fibroblasts to induce mouse mammary epithelial branching *in vitro* (Sumbal and Koledova, 2019). In addition, perturbed hedgehog signaling (Zhao et al., 2017) or integrin activity regulation (Peuhu et al., 2017) in the mouse mammary gland stroma was shown to delay pubertal mammary gland branching morphogenesis and to reduce the regenerative capacity of mammary epithelium. In the light of these previous data, it is plausible that elevated amounts of intralobular fibroblasts in highly branched, more proliferative, TDLUs could be functionally linked to the elevated branch count. However, such causality remains to be confirmed.

In earlier studies conducted with lower resolution imaging, TDLUs were divided into four different types of branching, where type I represents young nulliparous tissue (on average 11 acini/TDLU), type II mature nulliparous tissue (on average 47 acini/TDLU), type III parous tissue (on average 81 acini/TDLU), and type IV intense branching occurring only during pregnancy and lactation (Russo et al., 1992). Of the TDLU samples analyzed quantitatively in this study, only a small fraction contained more than 81 branches in total, thus reaching category III suggested to predominate in parous women (Russo et al., 1992). Importantly, our data suggest that the TDLUs of young nulliparous and mature parous donors are, in fact, not significantly different in branch count. Similar conclusions have also been made by others who have investigated human TDLUs with tissue sections, and concluded that the length of the postpartum period is an important factor affecting branching (Jindal et al., 2014; Ogony et al., 2022). In our sample cohort the shortest known postpartum period to sample collection was 6 years. Although the dataset analyzed here is still relatively small and lacks the ability to distinguish the effect of age from parity, our data suggest that the effect of parity on TDLU branching may not be as straightforward as previously described.

In addition to parity and age, menstrual cycling (Ramakrishnan et al., 2002) and hormonal contraception (Russo et al., 1992) have been indicated to affect TDLU branching. The samples analyzed here represented different menstrual cycle phases or hormonal contraception. While no differences in the TDLU branch count could be clearly linked to the hormonal contraception status, an elevated branch count was detected in the samples from donors in mid cycle, corroborating earlier findings demonstrating increased proliferation at a similar phase (Going et al., 1988; Söderqvist et al., 1997; Ramakrishnan et al., 2002). This increase is followed by a decrease towards the end of the cycle, which matches reported increase in apoptosis (Anderson et al., 1982; Longacre and Bartow, 1986). Further, the position of the TDLU within the whole gland could affect the branching type, as fibroglandular density, along with tumor incidence, has been indicated highest at the upper outer quadrant of the breast in Western women (Chen et al., 2017). The samples in our branch count analysis were collected from reduction mammoplasty operations, where the tissue is resected from the lower quadrants of the breast.

Higher ER expression was observed in the TDLU epithelium of samples from mature parous women compared to young nulliparous. Gradually increasing ER status has been previously reported in response to ageing (Santandrea et al., 2021; Shoker et al., 1999) and ER+ breast cancer is more common with increasing age (Shah et al., 2022). In turn, parity and the use of hormonal contraception have been shown to decrease ER levels (Russo et al., 1999; Williams et al., 1991; Hilton et al., 2018). In our sample cohort where the effects of age and parity cannot be separated, the effect of ageing appears to dominate over parity. Also, T-cell infiltration into mammary epithelium was more prominent in TDLUs of mature parous donors. Similar effects of age or parity have not been previously reported and a more detailed characterization of this T-cell population is warranted.

Overall, this study presents the first systematic analysis and mathematical modelling of the glandular 3D structure in normal human breast epithelium. Our findings add a new layer of detail on the anatomy of the human breast. We demonstrate that 3D histology or volumetric imaging of optically cleared entire TDLUs can be used for studying the relationship between TDLU architecture and the cell types that form it. This study also provides a reference data set for future development of light sheet image analysis, and a protocol and framework for future investigations on the role of ductal architecture in the regulation of growth in healthy and malignant breast epithelium, including carcinogenesis and invasive progression. Importantly, the identities and niches of distinct cell types within the TDLU structure are likely to become clearer as research proceeds, revealing previously unknown structure-function relationships in the breast tissue.

## Materials and methods

### Human breast tissue

At the Turku University Hospital, normal breast tissue was obtained from donors undergoing breast reduction mammoplasty. Normal breast tissue adjacent to the tumor was collected in mastectomy due to pre-invasive carcinoma. Tissues were voluntarily donated upon written informed consent at Turku University Hospital (Ethical approval ETKM 23/2018), and the donors filled a questionnaire providing details of the reproductive history and hormonal medication. At Mediclinic in Oud-Heverlee (BE), human breast tissues were anonymously retrieved from healthy donors who underwent voluntary cosmetic reduction surgery in the absence of any medical indication. All donors were adequately informed by third party clinicians about the use of the tissue and provided written informed consent for the use of the resection material and publication of the experimental results. Limited and fully anonymous donor information (age, parity status, familial cancer risk) was obtained without any link to the donor’s medical record. All tissues were surgically dissected to obtain pieces with glandular tissue, and processed for fixation. For samples with high apparent mammographic density, a mild pre-digestion in collagenase was utilized before fixation (1.5h with 300 U/ml collagenase [Sigma, C7657]).

### FUnGI clearing

Tissue clearing and immunolabelling of tissue pieces was done following previously described FunGI protocol (Rios et al., 2019). All incubation steps were performed on a roller mixer. Tissue pieces were fixed in 4% PFA in +4°C for 24 h, and stored in PBS + 1:100 penicillin/streptomycin at +4°C until labelling. Fixed tissue pieces were labelled with DAPI (1:1000-1:3000) in PBS for 2 days in +4°C. The areas with TDLU structures were visualized with fluorescence microscopy, and chosen areas cut into pieces of approximately 2-5 mm in diameter. Tissues were washed in wash buffer [WB; 50 µg/ml ascorbic acid (Sigma, A4544), 0.05 µg/ml L-glutathione (Alfa Aesar, J62166) in PBS + 0.1% Tween] for 30 min, and in WB1 [50 µg/ml ascorbic acid, 0.05 µg/ml L-glutathione in PBS + 0.2% Tween + 0.2% Triton X-100 (Sigma, T8787) + 0.2% SDS (Amresco, 0227) + 0.2% BSA (Biowest, P6154)] for 2-3 hours in +4°C, and incubated in anti-keratin 8 primary antibody [1:100 (Hybridoma Bank, TROMA-1)] diluted in WB2 (50 µg/ml ascorbic acid, 0.05 µg/ml L-glutathione in PBS + 0.1 % Triton X-100 + 0.02% SDS + 0.2% BSA) for 2 days in +4°C. After, pieces were washed 3×1 h in WB2, and incubated in anti-rat secondary antibody [1:400 (Thermo Fisher Scientific, Alexa Fluor 594)] diluted in WB2 for 36 h. Pieces were then washed again 3×1 h in WB2, after which they were moved to ∼25-30 ml of freshly prepared FUnGI solution [FUnGI base: 50% glycerol (Honeywell, 15675690), 2.5 M fructose (ACROS Organics, 1064642), 2.5 M urea (ACROS organics, 424585000), 10.6 mM Tris Base, 1 mM EDTA. Added prior to clearing: 50 µg/ml ascorbic acid, 0.05 µg/ml L-glutathione] and incubated for 2 hours in room temperature. Finally, pieces were moved to ∼25-30 ml of fresh FunGI solution and incubated at minimum o/n in +4°C, or until mounting.

### iFLASH clearing

Tissue clearing and immunolabelling were adapted from previously described protocol (Messal et al., 2021). All incubation steps were performed on a roller mixer. Tissue pieces were fixed with either 4% PFA in SDS permeabilization buffer (10% SDS in H_2_O, pH 7.4) o/n in +4°C or with 2%PFA in SDS permeabilization buffer o/n in RT. The next day, pieces were washed 3×5 min with PBS and moved to SDS permeabilization buffer for 1-2 days in +37°C. Fixed tissue pieces were then transferred to 25-50ml of CUBIC 1 [1.6 M urea, 5% Quadrol (Sigma, 122262), 15% Triton X-100, 25 mM NaCL (Fisher Scientific, S/3160/60) in ddH_2_O] (Susaki et al., 2014) and incubated in +37°C for a minimum of 3 weeks, changing the buffer every 2-3 days. Next, the pieces were labelled with DAPI [1:1000-1:3000, (Life Technologies, D1306)] in PBS o/n at room temperature. The areas with TDLU structures were visualized with fluorescence microscopy, and chosen areas cut into pieces of approximately 2-5 mm in diameter. Tissues were then permeabilized again using SDS permeabilization buffer for 2 days in

+37°C, changing the buffer between the days. The tissues were then washed 3×1h in PBT (PBS + 0.2% Triton X-100, pH 7.4) at room temperature and blocked with iFLASH blocking buffer [10% FBS (Sigma, 122262), 5% DMSO (Chem Cruz, 358801), 0.1% NaN_3_, 1% BSA in PBT] for 1h at room temperature, and incubated in primary antibodies diluted in iFLASH blocking buffer for 3 days [anti-keratin 8 primary antibody 1:100 (Hybridoma Bank, TROMA-1); anti-keratin 14 primary antibody 1:100 (Biolegend, 905301)]. Tissues were washed 3×1h with PBT at room temperature, and pieces incubated in secondary antibodies [anti-rat secondary antibody (H+L) 1:400 (Thermo Scientific, Alexa Fluor 488), anti-rabbit 568 secondary antibody (H+L) 1:400 (Thermo Scientific, Alexa Fluor 568)] in +37°C. Finally, the pieces were washed 3×1h with PBT at room temperature, and transferred to 25-50ml of CUBIC2 [1.2 M sucrose (Millipore, 107651), 3.6M urea, 9% triethanolamine (Sigma, 90279), 0.1% Triton X-100 in ddH_2_O] (Susaki et al., 2014) for a minimum of 2 days in +37°C, or until mounting and imaging.

### Tissue sample and bead mounting

Labelled and cleared tissue pieces were imaged again to trim and position them optimally for the light sheet imaging excitation and emission light paths. Pieces were mounted in 3.5 mm dishes in 2% agarose (Lonza, 50180) diluted in milliQ water (FunGI protocol) or CUBIC2 (iFLASH protocol), and let set at +4°C. The solidified agarose blocks were cut out and trimmed to approximately 8 mm in diameter to fit available imaging set up. Blocks were transferred to 25-30 ml fresh FunGI or CUBIC2 respectively and incubated at minimum o/n in +4°C, or until imaging.

For deconvolution, bead samples were prepared for point spread functions. Beads (Fluoro-max Fluorecent Polymer Microspheres, Thermo Scientific) were diluted 1:10 000 - 1:20 000 in 2% agarose (as described previously) and vortexed well for uniform distribution. Mixture was let settle for 10 min at room temperature to allow bubbles to rise to the surface, and then mounted into an Eppendorf lid. The solidified pieces were then cut out. Additionally for FunGI protocol, bead samples were incubated in FunGI o/n before imaging.

### Immunohistochemistry of frozen tissue sections

Tissue samples were fixed overnight at +4°C in periodate–lysine–paraformaldehyde (PLP) buffer [1% paraformaldehyde (Thermo Scientific, 28908), 0.01M sodium periodate (Sigma, 311448), 0.075 M L-lysine (Alfa Aesar, 15404639) and 0.0375 M P-buffer (0.081 M Na2HPO4 and 0.019M NaH2PO4; pH 7.4)]. After washing twice with P-buffer, samples were incubated in 30% sucrose in P-buffer for a minimum of two days. Samples were mounted in Tissue-Tek® O.C.T. Compound (Sakura, 4583) on dry ice and cut into 8 μm sections.

For immunofluorescence labelling, the frozen sections were first thawed for 1 h at room temperature. The sections were blocked and permeabilized in permeabilization buffer (2% BSA, 0.1% Triton X-100 in PBS) for 30 min at room temperature. Primary antibodies were incubated in 2% BSA in PBS overnight at +4°C [anti-keratin 8 primary antibody 1:1000 (Hybridoma Bank, TROMA-1); anti-keratin 14 primary antibody 1:1000 (Biolegend, 905301)]. Then, the sections were washed 3 x 10 min with PBS and incubated with the secondary antibodies, diluted in 2% BSA in PBS, for 1 h at room temperature [anti-rabbit secondary antibody (H+L) 1:400 (Thermo Scientific, Alexa Fluor 488), anti-rat 568 secondary antibody (H+L) 1:400 (Thermo Scientific, Alexa Fluor 568)]. Sections were then washed 3 x 10 min with PBS (1:1000 DAPI in the second wash) and then 5 min with milliQ water. Finally, the sections were mounted under a #1.5 glass coverslip with Mowiol® (Calbiochem) supplemented with 2.5% 1,4-diazabicyclo[2.2.2]octane (DABCO, Sigma-Aldrich).

### Light sheet microscopy and deconvolution

Samples were imaged with MSquared Aurora Airy Beam Light Sheet microscope using Aurora acquisition software version 0.5. The objective used was a Special Optics dipping objective which has dipping medium refractive index (RI) dependent magnification 15.3×-17.9× (NA 0.37-0.43) in immersion medium RI range 1.33-1.56. Alexa488 secondary antibody was imaged with a 488 nm laser and emission filter at 500-540 nm and Alexa594/Alexa568 secondary antibodies with a 568 nm laser with long pass emission filter at 570 nm. 3D images were acquired by taking z-stacks with 400 nm spacing with a pixel size of 387 nm. Images were deconvoluted using Aurora Deconvolution software version 0.5 using point spread functions of fluorescent beads obtained from agarose embedded samples.

### Confocal imaging

Cleared tissue samples were placed in 3.5mm glass-bottom dishes (Cellvis, D35-14-1-N) for imaging, additionally under a #1.5 coverslip covered in Vectashield (Bionordika, H-1000-10) if necessary to reach maximum imaging depth. Samples were imaged with the 3i Marianas CSU-W1 spinning disk (50µm pinholes) confocal microscope using SlideBook 6 software. The objective used was a 10x Zeiss Plan-Apochromat objective (NA 0.45). Signal was detected with the following solid-state lasers and emission filters; 405nm laser, 445/45nm; 488nm laser, 525/30nm; 561nm laser, 617/73nm. 3D images were acquired by taking z-stacks with a 2µm step size using a Photometrics Prime BSI sCMOS camera with a pixel size of 6.5 μm. Images were 16bit with pixel dimensions 2048×2048.

Tissue sections labelled with immunofluorescence were imaged at the same setup using the 20x Zeiss Plan-Apochromat (NA 0.8) and 63× Zeiss Plan-Apochromat (NA 1.4) objectives. Images were acquired by taking a z-stack with 6 slices with the step size of 1 μm.

### Image segmentation and branching analysis

Only images reliably showing a substantial fraction of the TDLU volume were chosen for images analysis (n=14). Segmentation of the 3D image was done using the Trainable WEKA Segmentation plugin (Arganda-Carreras et al., 2017). Images were preprocessed with background subtraction by rolling ball algorithm. A random forest pixel classifier was trained individually to each 3D image. 3-5 of the images in each image stack were used as training data, and a 200 tree and 2 feature random forest classifier was trained using Gaussian blur, mean, maximum, minimum, median, Laplacian, entropy, Sobel, difference of Gaussians, bilateral and Kuwahara filters as features. Segmentations were manually checked and fixed for mistakes.

### Quantitative analysis of the segmented TDLU structures was done by FilamentTracer function of Imaris

8.1.2 and 10.0.0 (Bitplane ltd.). After automatic placement of seed points using a minimum size of 25 µm estimated from the minimum branch diameter in the data, seed points were manually checked and edited when necessary. Model surface was manually thresholded to match the surface of the TDLU branches in the data. Branch diameters were calculated as the shortest distance from the distance map. Resulting filament models were extensively manually checked for errors and corrected before quantitative measurements were exported. Of all the analyzed TDLUs only the ones showing the whole TDLU volume, and in which the connectedness of the branches could be reliably traced were selected for analyses concerning topological properties of the whole TDLU (n=8). Filament models of TDLU structures were exported as TIFF stacks from Imaris and further analyzed using Tree Surveyor version 1.0.12.1 (Short et al., 2013) to extract parameters of subtree ancestry, network connectedness and the branching rotation angles. Rotation angles where extracted as the sharp (0-90⁰) angle between two planes defined by branches in two successive branching events.

Localization of KRT14^hi^ cells was observed from analyzed filament models in Imaris, and matched to corresponding subtree ancestry and network connectedness in Tree Surveyor. Total branch count of each TDLU was extracted from finalised filament models. TDLU volume was estimated by perpendicularly measuring the length and width of each structure and assuming a regular shape applying the formula for an ellipsoid’s volume;

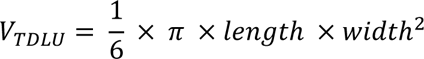

### Paraffin embedding of cleared tissue samples

After light sheet imaging, cleared tissue pieces were stored in CUBIC2 in +4°C for up to 4 months. For re-embedding into paraffin, pieces were manually extracted from the agarose mounting and remounted in HistoGel (Epredia™, HG-4000-012). HistoGel-mounted tissue pieces were re-embedded into paraffin blocks, from which a 3mm core containing the tissue of interest was taken, moved to a new mold, melted down and re-embedded into paraffin in a TMA formation. The constructed TMA blocks were then serial sectioned with a 4µm section thickness up to 1.5cm of depth, or until tissue was finished. Every 10^th^ section was then H&E stained for 3D reconstruction, and remaining sections used for Opal 7-plex immunolabelling at planes of interest.

### 3D reconstruction

The 3D reconstruction for the TMA spot selected for visualization was done following a pipeline modified from (Ruusuvuori et al., 2022). First, the stack was non-linearly aligned using the TrackEM2 plugin in Fiji/ImageJ v2.14.0 (Schindelin et al., 2012). Following registration, acini were manually segmented in Qupath v0.5.1 using the brush tool (Bankhead et al., 2017). The acquired masks were then employed to discard background regions from the images in Python. Finally, the 3D reconstruction was created in Matlab R2019a using the Vol3D v2 script (Woodford, 2024).

### Multiplex staining

Multiplex immunohistochemistry (mIHC) staining was performed using the BOND RX automated stainer (Leica Microsystems Ltd). For the multiplex mIHC we used the Opal 6 plex detection kit (Akoya Biosciences, NEL871001KT) and staining was conducted according to the manufacturer’s protocol. All reagents were prepared in advance and slides were loaded on the instrument. To begin the staining process, PFA-fixed and paraffin embedded tissue slides (5 µm) were incubated for 1h at 60°C with Bond Dewax solution (Leica Bond, AR9222) to deparaffinize tissue sections. Antigen retrieval was achieved by immersing the sections with BOND Epitope Retrieval Solution 1 (Leica Bond, AR9961) for 20 minutes at 95°C. Subsequent blocking of non-specific binding sites was followed for 15 minutes at RT. According to the primary Ab species, a different blocking buffer was used (for mouse and rabbit primary Ab: Akoya Biosciences, ARD1001EA; for rat primary Ab: Akoya Bioscience, 30024). Next, slides were incubated for 30 minutes at RT with primary Ab which was diluted in TNB buffer (Akoya Biosciences, FP1020), followed by incubation with secondary antibodies ImmPRESS-HRP anti mouse/rabbit (Akoya Biosciences, ARH1001EA) or ImmPRESS-anti rat HRP (Akoya Biosciences, 30032) for 10 minutes at RT. Following this step, an Opal fluorophore working solution (Akoya Biosciences, NEL871001KT) was applied for 10 minutes at RT. All Opal reagents were diluted 1:150 in an amplification buffer (Akoya Biosciences, FP1609) except Opal 780, which was diluted to 1:50 in TNB buffer. To finish the cycle, primary and secondary Abs were stripped with BOND Epitope Retrieval Solution 1 for 20 minutes at 95°C. Subsequent staining rounds followed the same steps. Finally, nuclei were counterstained with DAPI (Akoya Biosciences, FP1490) for 5 minutes at RT before mounting the stained tissue with mounting medium (Invitrogen, P36961). **Table S3** provides detailed information on the primary Abs used. Stained sections were imaged using a slide scanner (Akoya vector Polaris).

### Multiplex image analysis

The branch counts of singular TDLUs were determined from consecutive H&E sections by adding up all distinguishable acini per structure. Matching TDLUs were then located in multiplex imaging and cropped out for individual analysis. Then, relevant epithelial area was segmented; ER to luminal epithelium (KRT19), p63 to basal epithelium (αSMA), Ki67 to epithelium (pan-keratin), CD3 to within [epithelium] or outside of luminal epithelium [intra-TDLU stroma] (KRT8) and S100A4 to within [epithelium] or outside of epithelium [intra-TDLU stroma] (CDH1). The resulting mask was then applied to the signal of interest, from which either its area (p63, CD3, S100A4) or number of nuclei (ER, Ki67) were determined. For stromal signal analysis, intra-TDLU was determined by connecting peripheral acini, and then removing the signal of interest from within the epithelium.

### Multiplex image analysis

The branch counts of singular TDLUs were determined from consecutive H&E sections by adding up all distuinguishable acini per sturcture. Matching TDLUs were then located in multiplex imaging and cropped out for individual analysis. Then, relevant epithelial area was segmented; ER to luminal epithelium (KRT19), p63 to basal epithelium (αSMA), Ki67 to epithelium (pan-keratin), CD3 to within [epithelium] or outside of luminal epithelium [intra-TDLU stroma] (KRT8) and S100A4 to within [epithelium] or outside of epithelium [intra-TDLU stroma] (CDH1). The resulting mask was then applied to signal of interest, from which either its area (p63, CD3, S100A4) or number of nuclei (ER, Ki67) were determined. For stromal signal analysis, intra-TDLU was determined by connecting peripheral acini, and then removing the signal of interest from within the epithelium.

### Mathematical modelling

BARW simulations were implemented as in (Hannezo et al., 2017) including branch self-repulsion (Uçar et al., 2021). The simulated networks will grow indefinitely unless growth is somehow limited. We tested two mechanisms for network growth limiting: 1) Growth was limited to a predefined volume, and 2) a model in which branches have branching potential that wanes in older generations and eventually stops the branching. 1000 simulations were run for both models to generate the dataset.

#### 1) Volume-limited model

At each simulation step each active branch can do one of three possibilities: (1) it can grow by one more step, (2) it can bifurcate to two child branches or (3) it can terminate (i.e. stop growing and become inactive) if it meets an existing branch or limits of the simulation volume. Each branch has a propagation direction unit vector ***K̂***(*t*) pointing to the direction of branch growth at time *t*. At each simulation step a random number in [0,1] is drawn and bifurcation happens if the number is smaller than the bifurcation probability. Otherwise the branch grows forward one step in direction ***K̂***. Similar to (Hannezo et al., 2017), at each step the branching angle is diffused by a random angle in range [0, 15⁰] (angle range determined as in (Roos et al., 2022)) and a rotation angle in range [0, 360⁰] to prevent unnatural line-straight branches. If bifurcation happens, two bifurcation angles *β* are drawn from a Gaussian distribution fitted to experimental bifurcation angle distribution (**Fig. S3G**), and two angles *θ* are drawn from a uniform angle distribution of [0, 360⁰] to represent the rotational directions of the new branches around the parent branch (**Fig. S3F**). An active branch terminates if its tip gets within termination distance *r*_*T*_ from an existing branch network point. Simulation starts with one active tip starting to grow with ***k̂*** = ***ê_x_*** Simulation runs until there are no active tips left.

3D vector rotations are done by the Rodrigues’ vector rotation (Vince, 2021).

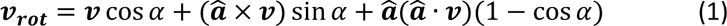

where *v* is the original vector to be rotated by angle *α* around the axis defined by the unit vector *α̂*, and *v*_rot_ is the final rotated vector.

Direction vectors ***K̂*** of the new branches in 3D space are calculated using the following algorithm: first a reference unit vector defining a direction perpendicular to parent branch direction is calculated as 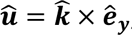, where *ê*_*y*_ is the standard unit vector along *y*-axis. Vector ***û*** is then rotated by the rotation angle *θ* around parent branch’s ***K̂*** vector using Eq. (1) to get a randomly in [0,360⁰] oriented rotation axis ***â*** for the final rotation with the branching angle *β*. The direction of the child branch is then calculated by rotating the parent branch’s direction vector ***K̂*** by branching angle *β* around the axis ***â*** again using Eq. (1).

Simulation parameter values were derived from the experimental data as follows and they were scaled to represent units of µm. Branch diameter value of 40 µm was used which was calculated by adding an extra 5 µm for the basal cell layer to the experimentally observed 35 µm branch diameter of the luminal cell layer. Bifurcation probability was calculated to be 0.15 by dividing the simulation step size 10 µm with the experimentally observed average branch length of 71 µm.

Branch repulsion causes an additional displacement to the direction vector ***K̂***. Every active tip experiences repulsion from all other existing network points that are within repulsion radius in the front half-space of the tip. Displacement due to repulsion 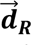 was calculated as 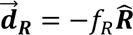, where *f*_*R*_ is the repulsion strength and _**R̂**_ unit vector pointing from the examined tip to the mass centre of all network points within repulsion radius in the fron half-space. 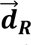 was added to ***K̂*** and the result was normalized to give direction unit vector for the next step.

Termination distance was determined from the experimental data by finding terminated branches that pointed towards another branch and measuring the gap from the tip of such branch to the surface of the second branch. The average gap in such cases was 18 µm (SD 11 µm) (**Fig. S6H**). Since the calculations in the model are done with branch paths that have thickness of zero (solid lines in **Fig. 5A**), the termination distance was set equal to 38 µm consisting of the gap plus half the branch diameter (**Fig. 5A**). To prevent immediate termination by the parent and sibling branch at the branching event, termination events only in the front half-space of the active tip were taken into account like in (Roos et al., 2022). As observed previously (Roos et al., 2022), simulation produces realistic networks only if immediate re-branching after a branching event is prevented and therefore a branching free zone of radius equal to branch diameter was established around each branching event to prevent immediate re-branching which sets a requirement that branches shorter than their width are not mature enough to branch again (**Fig. 5A**). Self-repulsion from closeby branches (Uçar et al., 2021) was used and the value of repulsion strength was optimized to 0.2 by finding the best match between the branch count distributions of the simulated and experimentally observed networks (**Fig. S7**). Repulsion distance was determined from the experimental data by finding branches that had been growing towards another branch but making a bend when getting close. The distance from the bend to the surface of the second branch was on average 35 µm (SD 11 µm) (**Fig. S6H**). The repulsion distance was set equal 55 µm consisting of the gap plus half the branch diameter again.

If a rigid predefined simulation volume box is used in the volume-limited model to limit the growth, some networks that start to grow at an angle with respect to the box can terminate prematurely by colliding to the sides of the simulation volume. There we did not confine the simulation to a predetermined box, but instead the network was allowed to grow in the direction it chooses. Simulation volume was controlled by terminating any branch that wanders further in length from the origin (maximum length) or wider from the general direction of the network (maximum width) than allowed by the limits (**Fig. 5B)**. General direction of the network was determined as a vector from origin to centroid of the set of current network points. Maximum length was set equal to experimentally observed average TDLU length of 460 µm and maximum width to 310 µm to create a limiting volume of typical TDLU size and having an aspect ratio of 1.5. In effect this creates a limiting volume that follows along the growing network.

#### *2)* Model of gradually waning branching potential

Mechanics of the simulation were otherwise identical to the volume-limited model, but in the branching potential waning model we modified the BARW mechanism so that when a branch stops growing and goes to the bifurcation state, instead of always bifurcating it makes a decision to bifurcate or self-terminate. Probability to self-terminate increases with branching level so that the probability to branch follows equation *p*_*B*_(*L*) = *q*^*L*^, where *L* is the branching level and *q* is a waning rate constant. This describes a fractional decline of branching potential by increasing branching level. Setting a requirement that *p*_*B*_(0) = 1, we estimated the waning rate *q* = 0.867 from a fit to the experimentally observed distribution of KRT14^hi^ cells (**Fig. 5C**) in different levels. In the simulation, when a branch reaches the end of elongation state (determined by the bifurcation probability parameter as in the volume-limited model), then a random number *R* in [0,1] is drawn, and if *R* < *q*^*L*^at the level *L* of the branch, it bifurcates as previously, but otherwise self-terminates and becomes permanently inactive.

Sensitivity analysis of the volume limiting model was done by varying one by one each model parameter by ±50 % around the value derived from experimental data. Effect of simulation limiting volume was investigated by varying the volume by ±20 % while retaining the volume aspect ratio of

1.5. 300 simulations were run on each tested parameter value and the results are summarized in **Fig. S5**. The model is most sensitive to the value of the branch diameter and branching probability. As expected, the simulation volume affects the size of the resulting network directly. Parameters for branch self-repulsion show the least effect on results. Reliability of the simulation is demonstrated by the fact that the best fit of simulation results to the experimental data is obtained by parameter values directly calculated from experimental data and the model depends weakly on the only parameter which could not be derived from the data, namely the repulsion strength.

The network topology characteristics depend strongly only on the decline rate parameter of the model of gradually waning branching potential (**Fig. S7**). Varying other parameters by ±50% produced mostly affected the physical size of the network which is seen as the varying branching density in **Fig. S7**. This model is very sensitive to the decline rate and it was only varied ±5% in the sensitivity analysis to produce reasonably comparable results.

Coefficient of determination *R^2^* was used to quantify the goodness of the model predictions *R*^2^ = 1 - 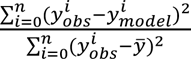, where 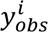 and 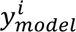 are the experimentally observed value and model prediction of data point *i*, and the average value of the observation calculated as 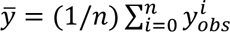 (Hannezo et al., 2017).

## Code availability

The mathematical model used for 3D simulation of TDLU branching is available on Mendeley Data (Peurla, 2023).

## Acknowledgements

We thank the patients that voluntarily donated their tissue to this study, the clinical staff of Turku University Hospital and Mediclinic, Oud-Heverlee (BE) for assisting in sample collection, Minna Santanen, Barbara Ramos Artigot, and Jukka Karhu for technical assistance, and all the lab members for constructive feedback over the course of this project. The Cell Imaging and Cytometry Core facility (Turku Bioscience, University of Turku, Åbo Akademi University, and Biocenter Finland), Biomedicum Imaging Unit (University of Helsinki), Histocore (Institute of Biomedicine, University of Turku) and Turku Bioimaging are acknowledged for services, instrumentation and expert advice. This work was supported by Sigrid Jusélius Foundation, Academy of Finland research fellowship (323096), and the Hospital District of Southwest Finland (11083). OP has been supported by Turku Doctoral Program of Molecular Medicine. LM would like to acknowledge support from Stichting Tegen Kanker (fellowship number ZKE4757).

## Author contributions

Conceptualization: OP, MP, EP; methodology: OP, MP, MV, LM, PR, CS, EP; formal analysis: OP, MP, LMK, JP, MV; investigation: OP, MP, LMK, JP, KS, ET, SS, MV, LM, NB, PB, PH; writing—original draft:

OP, MP, EP; writing—review and editing, all authors; visualization: OP, MP, MV; supervision: PR, CS, PH, EP; funding acquisition: CS, EP

## Supplemental information

**Figure S1.** Representative images of light sheet microscopy imaging of optically cleared human breast tissue related to Figure 1

**Figure S2.** Subtree structures with quantification of KRT14^hi^ cells related to Figure 2.

**Figure S3.** Light sheet microscopy imaging of transgelin and quantification of its localization within TDLUs, related to Figure 2.

**Figure S4.** The number of branches per TDLU in serially sectioned histological analysis graphed in relation donor age, related to Figure 3.

**Figure S5.** Representative images of HE-stained tissue sections of the donor samples used for quantitative branching analysis related to Figure 4.

**Figure S6.** Branching structure characteristics related to Figure 4.

**Figure S7.** Examples of subtree graphs of experimentally analyzed TDLUs and networks generated by both simulation models, related to Figure 5.

**Figure S8.** Sensitivity analysis of the volume-limited branching model, related to Figure 5.

**Figure S9.** Sensitivity analysis of the gradual waning branching potential model, related to Figure 5.

**Table S1.** Donor sample characteristics.

**Table S2.** Comparison of the simulation branching parameters.

**Video S1. TDLU 3D branched structure is revealed through the analysis of KRT8 signal.** Image volume sequence and Imaris filament tracer analysis of two adjacent TDLUs. KRT8 (red). Scale bar length shown (varies).

**Video S2. Visualization of iFLASH-cleared TDLU-structures reveals that KRT14^hi^cells localise mainly along the main subtree of the TDLU.** Image volume sequence, Imaris filament tracer analysis and surface rendering of a KRT14 (cyan) /KRT8 (red) labelled TDLU. Scale bar length shown (varies).

---

**Figure S1, related to Figure 1.**
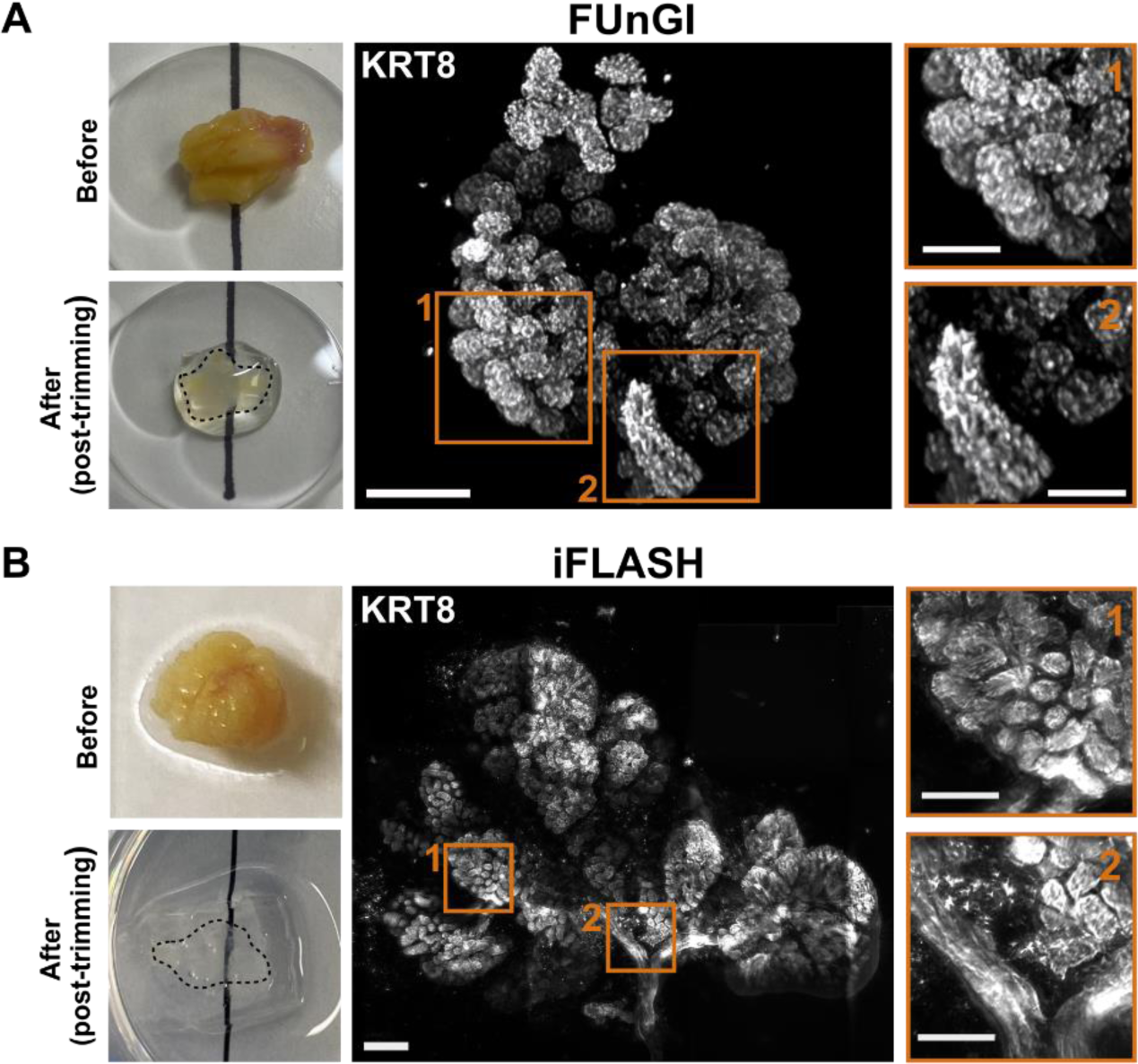
Light sheet microscopy imaging of optically cleared human breast tissue. Representative images of breast tissue before and after FunGI clearing (post-trimming) (**A**) or before and after (post-trimming) iFLASH clearing (**B**) are shown. Magnifications are shown for two ROIs. Scale bars: 200µm (main), 100µm (ROIs).

**Figure S2, related to Figure 2.**
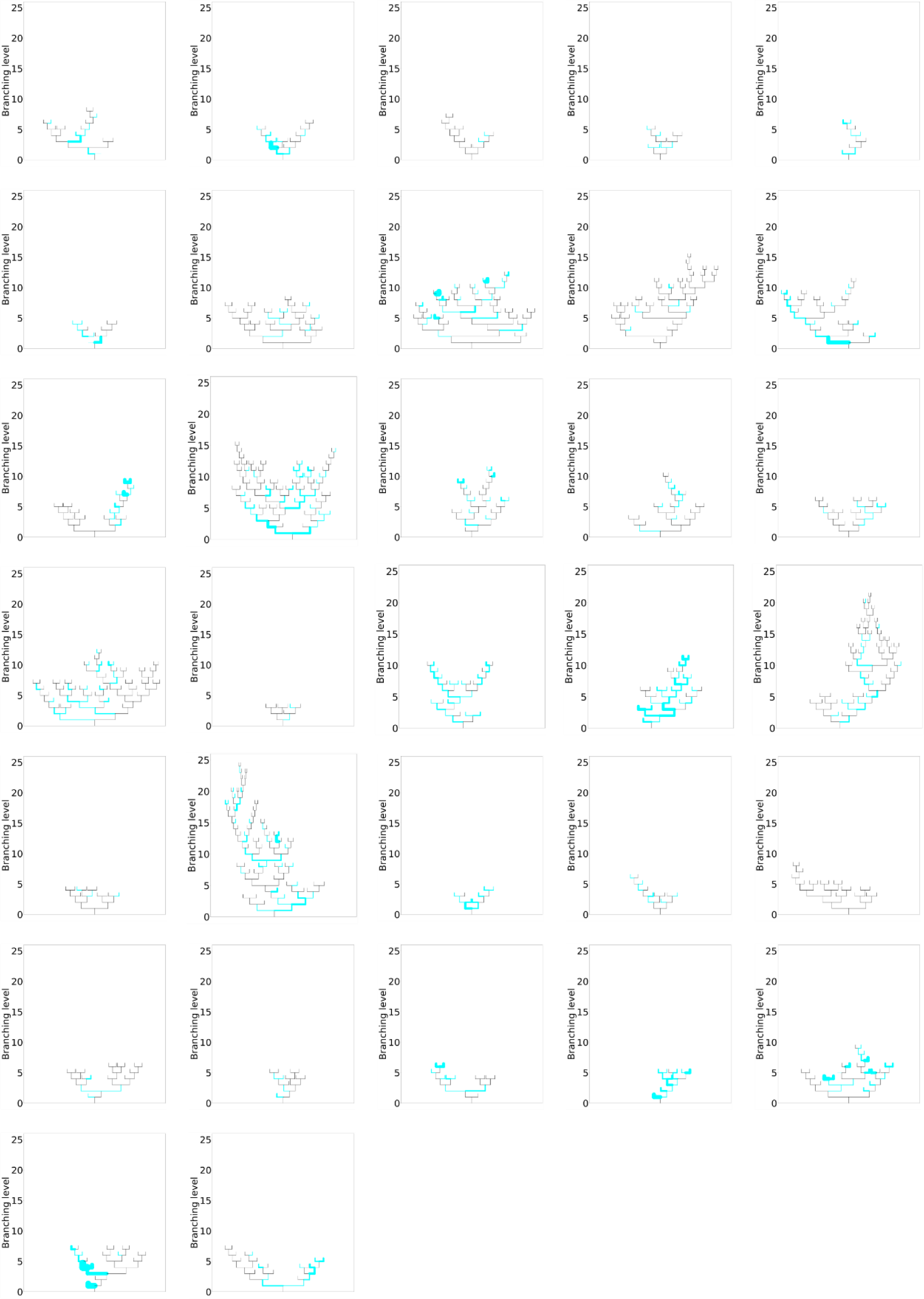
Subtree structures with quantification of KRT14^hi^ cells. Tree structure of TDLUs with the distribution of KRT14^hi^ cells within the branching hierarchy (cyan; line width is representative of number of KRT14^hi^ cells in each branch. Branch length not to scale.)

**Figure S3, related to Figure 2.**
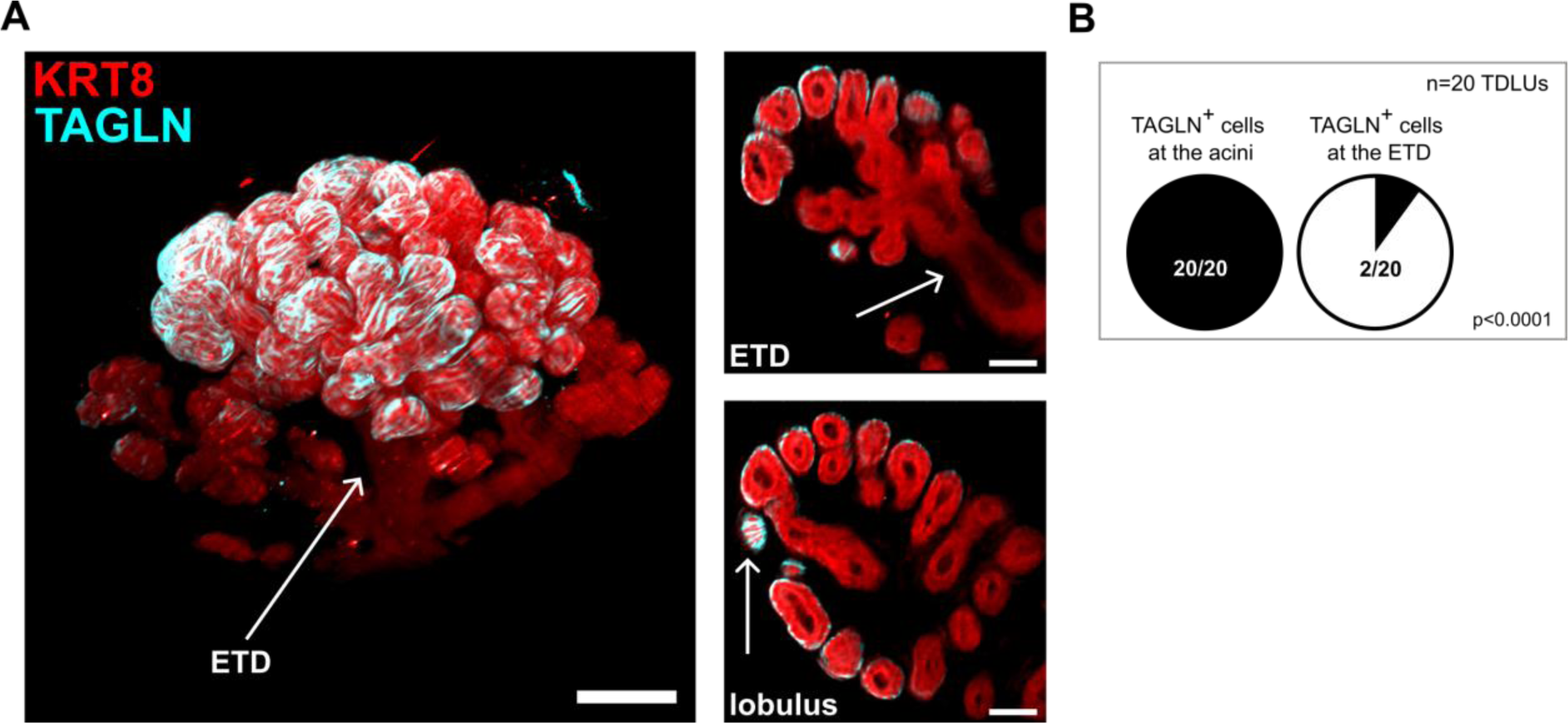
Light sheet microscopy imaging of transgelin (TAGLN in basal epithelial cells, cyan) and KRT8 (luminal epithelial, red) in entire TDLUs within cleared breast tissue pieces. TAGLN-positive cells are less visible in the single z-sections of ETD (ROI above) compared to the lobulus (ROI below) (**A**). Scale bars: 100µm (main), 50µm (ROIs). Quantification of TAGLN^+^ cell localization (n=20, Fisher’s exact test) (**B**). Images and quantification are representative of 20 TDLUs from 6 donors.

**Figure S4, related to Figure 3.**
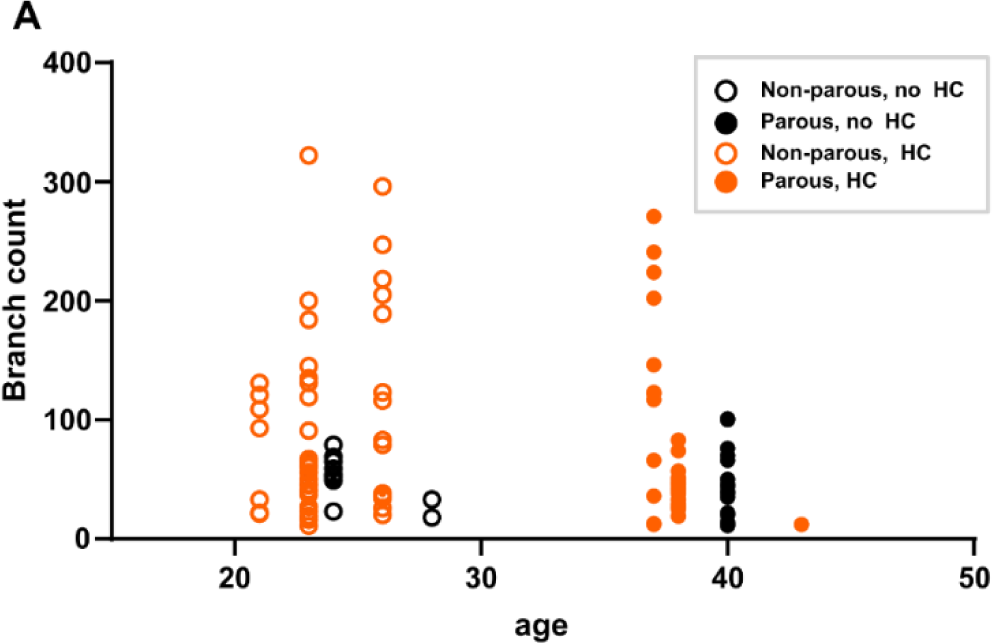
The number of branches per TDLU in serially sectioned histological analysis graphed in relation donor age. Parity of the tissue donor or the use of hormonal contraceptives is indicated (n=110 TDLUs from 10 donors).

**Figure S5, related to Figure 4.**
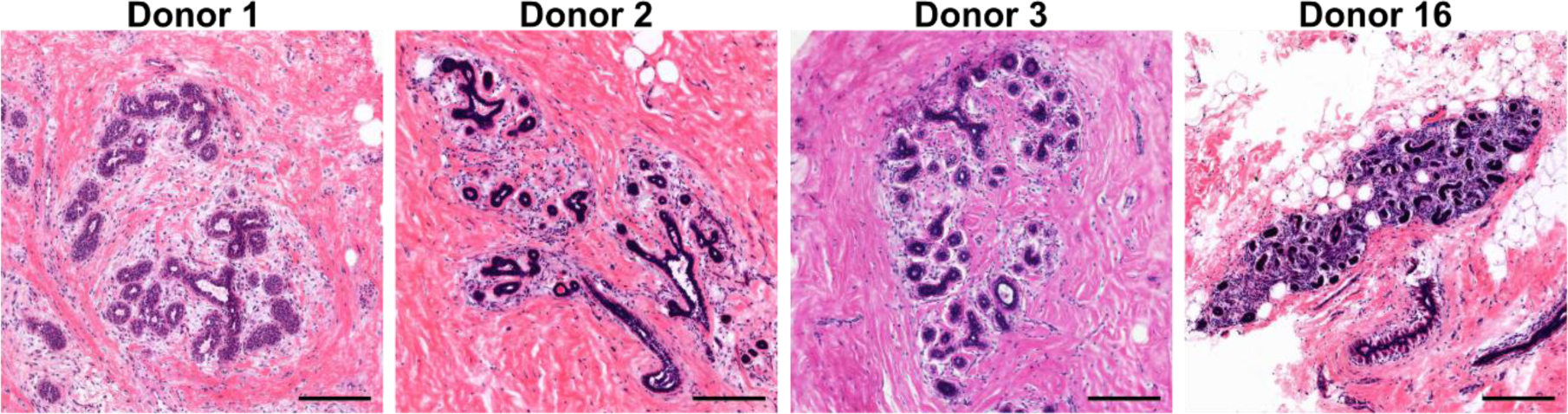
Representative images of H&E-stained TDLUs in breast tissue sections of the donor samples used for quantitative branching analysis in Figure 4. **Scale bars: 100µm.**

**Figure S6, related to Figure 4.**
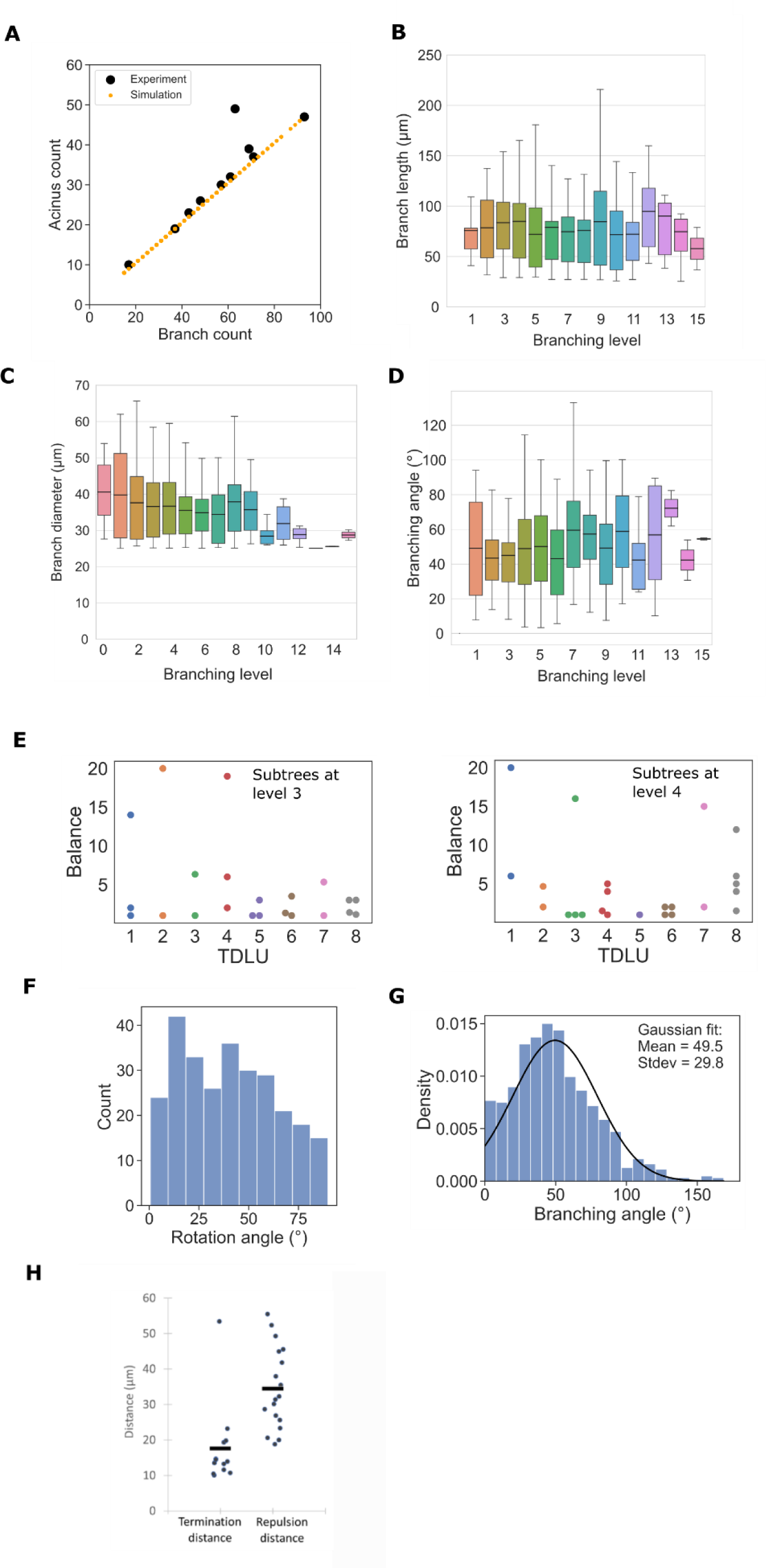
Branching structure characteristics. **A.** Total number of branches and number of acini in a TDLU and simulated branching networks. **B-D.** Experimentally observed branch length (**B**), branch diameter (**C**), and branching angle (**D**) at different branching levels (n_length_=670, n_diameter_=670, n_angle_=670 branches from 14 TDLUs of 4 donors). Boxes show median and interquartile range, and whiskers the full range of data points. **E.** The symmetry of each pair of subtrees originating from the same parent branch. Balance was calculated as the ratio of the number of network endpoints in each of the subtrees in the pair (Lefevre et al., 2017). Balance is shown for subtrees originating from levels 3 and 4 to rule out bias caused by selection of subtree starting level (n=8 from 4 donors). **F.** Distribution of the branching rotation angle. Distribution was approximated to be uniform in simulations (n=14 TDLUs of 4 donors). **G.** Distribution of the branching angle (branch opening angle). Fit shows the normal distribution used in the simulations to draw the branching opening angles (n=14 TDLUs of 4 donors). **H**. Distributions of termination and repulsion distances measured from the experimental data. Horizontal lines mark the average values. n=8 TDLUs of 4 donors.

**Figure S7, related to Figure 5.**
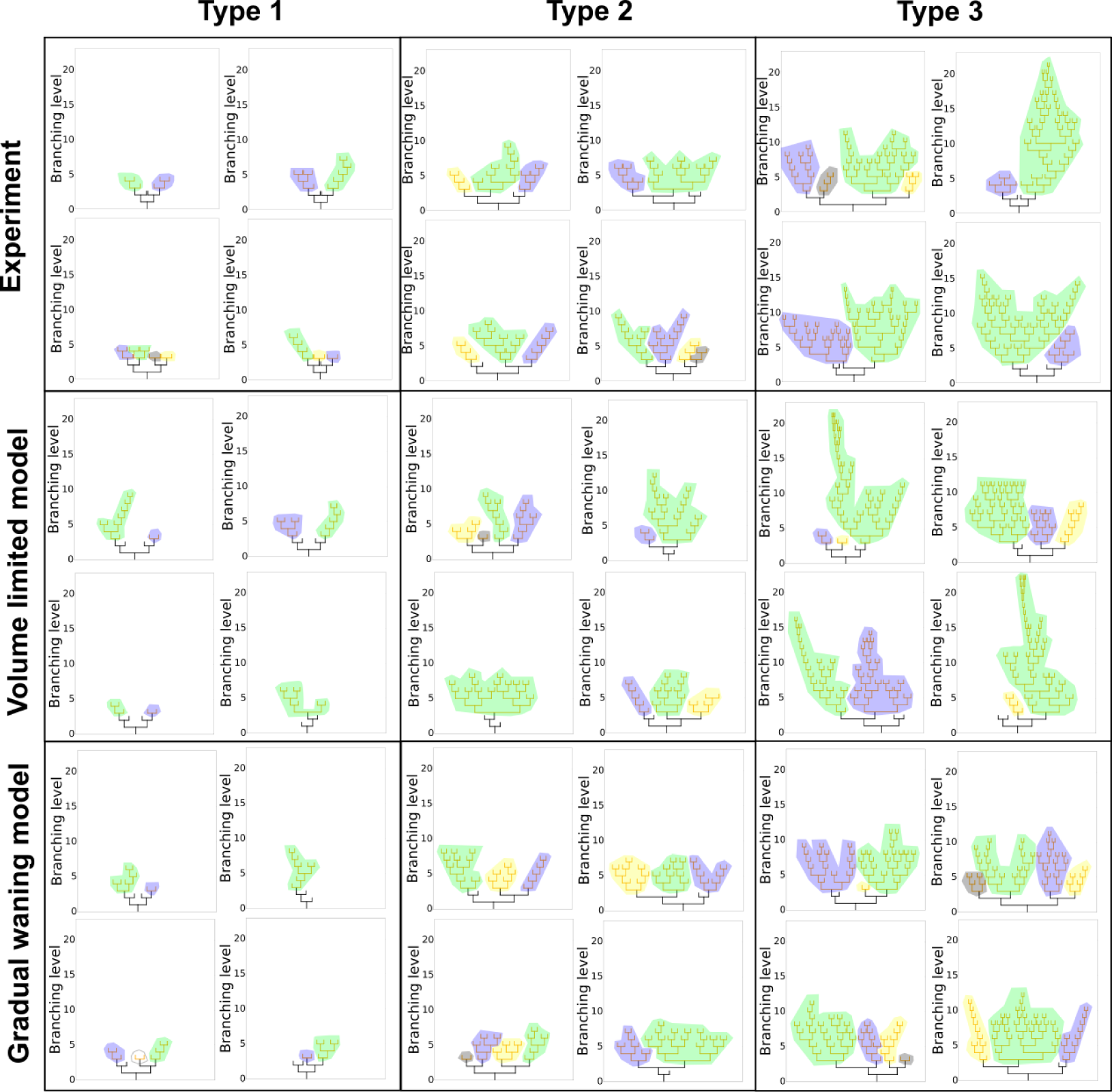
Examples of subtree graphs of experimentally analyzed TDLUs and networks generated by both simulation models. Trees having branch counts in the range of lobulus types 1, 2 and 3 (as shown in Fig1.) are shown and subtrees are highlighted by different colors for clarity.

**Figure S8, related to Figure 5.**
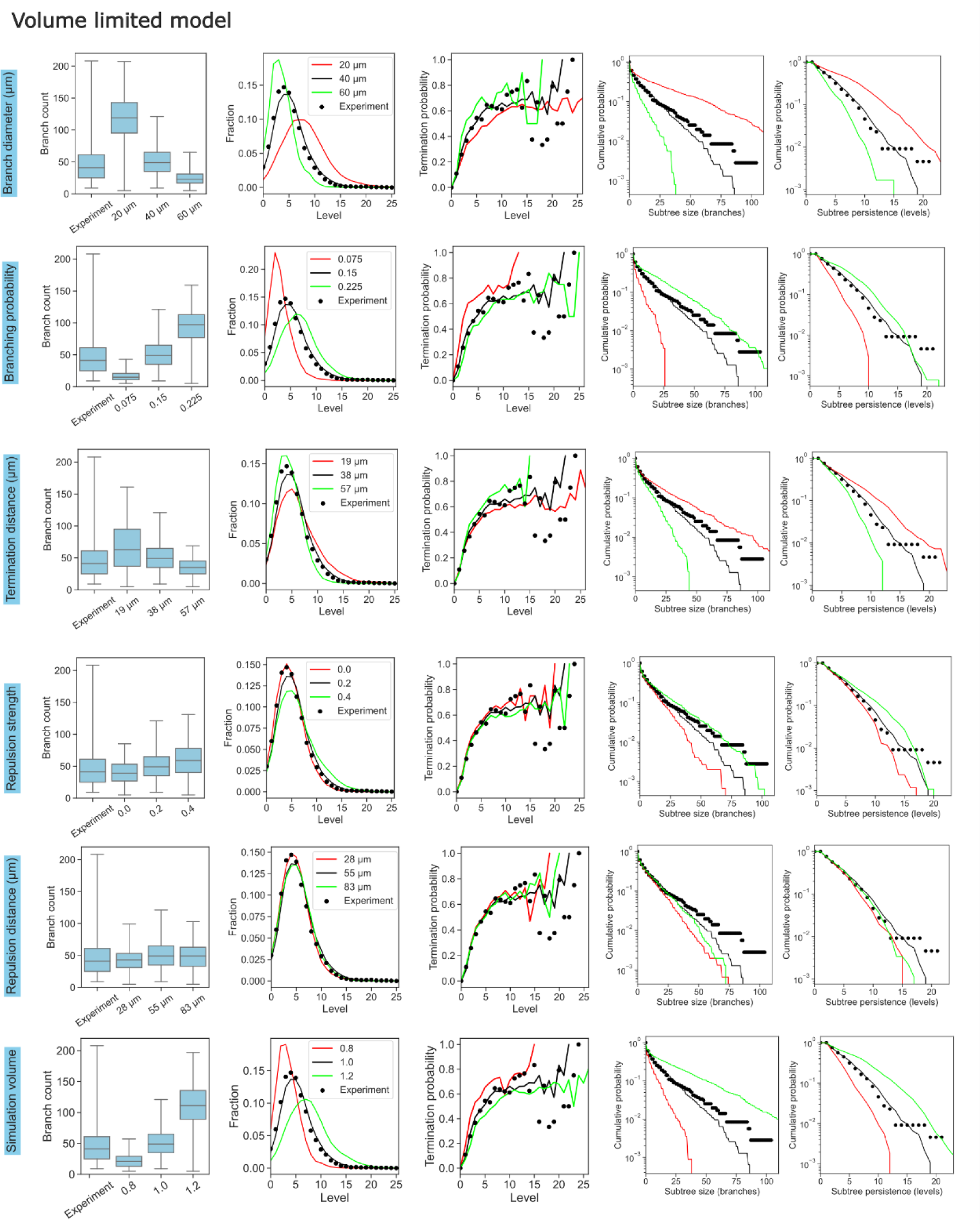
Sensitivity analysis of the volume-limited branching model. Simulation volume was varied by ±20 % and all other parameters ± 50% from the value determined from the experimental data. 300 simulations were run on each parameter value.

**Figure S9, related to Figure 5.**
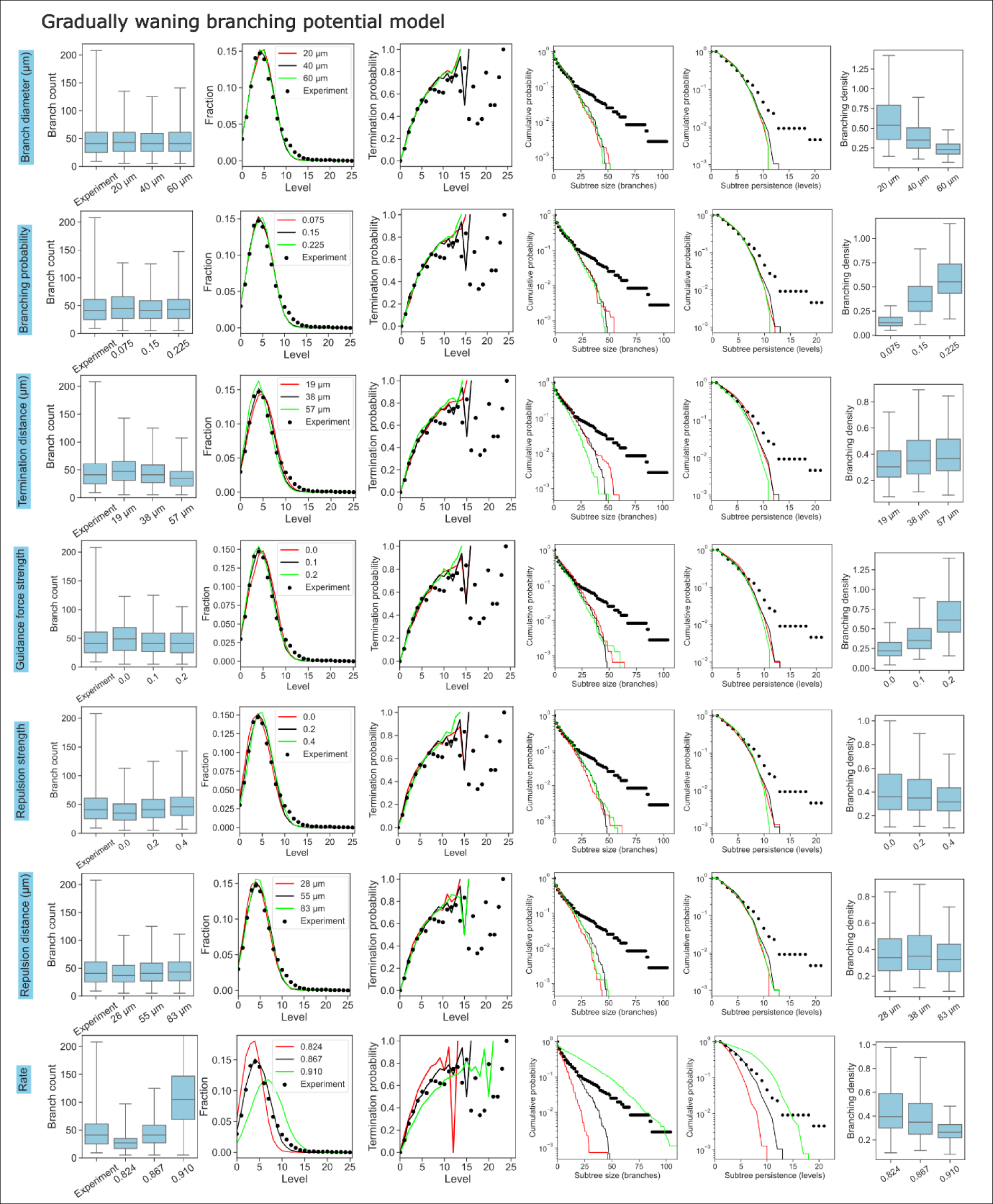
Sensitivity analysis of the gradual waning branching potential model. Decline rate was varied ±5% and all other parameters ± 50% from the value determined from the experimental data. 300 simulations were run on each parameter value.

**Table S1.** Breast sample donor characteristics. HC, hormonal contraceptives; IUD, intra-uterine device; N/A, not applicable; IHC, immunohistochemistry; BMI, Body mass index; DCIS, ductal carcinoma *in situ*.

| Donor | Age (years) | Parity | Procedure | Days since menstruation | HC type | Time since last pregnancy (years) | BMI | Clearing method | Branch length (μm) | Branch diameter (μm) | Branch angle (°) | Material notes |
| --- | --- | --- | --- | --- | --- | --- | --- | --- | --- | --- | --- | --- |
| *Turku University Hospital* |  |  |  |  |  |  |  |  |  |  |  |  |
| 1 | 24 | 0 | Reduction mammoplasty | N/A | oral | N/A | 29.07 | FUnGI | 74.1±40.4 | 34.7±9.5 | 49.9±30.6 |  |
| 2 | 43 | 2 | Reduction mammoplasty | N/A | IUD | 7 | 25.78 | FUnGI | 95.7±58.5 | 43.9±9.9 | 38.2±22.2 | Mastopathy |
| 4 | 18 | 0 | Reduction mammoplasty | 28 | none | N/A | 21.46 | iFLASH |  |  |  |  |
| 5 | 40 | 1 | Reduction mammoplasty | 14 | none | 8 | 26.86 | iFLASH, IHC |  |  |  |  |
| 6 | 21 | 0 | Reduction mammoplasty | N/A | implant | N/A | 28.41 | iFLASH |  |  |  |  |
| 7 | 37 | 2 | Reduction mammoplasty | N/A | IUD | 11 | 27.85 | iFLASH |  |  |  |  |
| 8 | 23 | 0 | Reduction mammoplasty | N/A | oral | N/A | 23.37 | iFLASH |  |  |  |  |
| 9 | 24 | 0 | Reduction mammoplasty | 19 | none | N/A | 24.13 | iFLASH |  |  |  |  |
| 10 | 40 | 1 | Reduction mammoplasty | 4 | none | 6 | 24.92 | iFLASH |  |  |  |  |
| 11 | 26 | 0 | Reduction mammoplasty | N/A | IUD | N/A | 24.77 | iFLASH |  |  |  |  |
| 12 | 42 | 1 | Reduction mammoplasty | N/A | IUD | 12 | 27.78 | iFLASH |  |  |  |  |
| 16 | 50 | 0 | Mastectomy | N/A | IUD | N/A | 29.74 | FUnGI | 88.8±50.5 | 34.0±5.9 | 49.6±29.9 | DCIS-adjacent tissue |
| 17 | 35 | 2 | Reduction mammoplasty | N/A | none | unknown | 26.99 | N/A (IHC) |  |  |  |  |
| 18 | 44 | 2 | Reduction mammoplasty | N/A | oral | unknown | 26.13 | N/A (IHC) |  |  |  |  |
| *Mediclinic, Belgium* |  |  |  |  |  |  |  |  |  |  |  |  |
| 3 | 48 | 2 | Reduction mammoplasty | 5 | none | unknown | unknown | FUnGI | 60.8±26.5 | 30.7±6.8 | 51.5±29.1 | 1.5h mild digestion |
| 13 | 38 | 4 | Reduction mammoplasty | N/A | oral | unknown | unknown | iFLASH |  |  |  |  |
| 14 | 28 | 0 | Reduction mammoplasty | 4 | none | N/A | 28.73 | iFLASH |  |  |  |  |
| 15 | 43 | 1 | Reduction mammoplasty | N/A | oral | unknown | 24.57 | iFLASH |  |  |  |  |

**Table S2.** Comparison of the simulation branching parameters.

|  | **Branch diameter** | **Branching probability** | **Branching angle** | **Simulation volume** |
| --- | --- | --- | --- | --- |
| **Human TDLU** | 40 $\mu\text{m}$ | 0.15 | 49° | 460×310×310 $\mu\text{m}^3$ |
| **Mouse mammary gland**<br>(Hannezo et al., 2017) | 3 a.u. | 0.10 | 30° | 280×150 a.u.<sup>2</sup> |

**Table S3.** Primary antibodies used for multiplex immunohistochemistry.

| Primary Abs | Company and catalogue number | Dilution | Species |
| --- | --- | --- | --- |
| P63 | Santa Cruz #sc-25268 | 1:500 | Mouse |
| Estrogen receptor | Cell signaling #13258S | 1:50 | Rabbit |
| Cytokeratin 19 | Invitrogen #MA5-12663 | 1:1000 | Mouse |
| FSP1 | Sigma-Aldrich #SAB5701199 | 1:1200 | Rabbit |
| Ecad | Abcam #76319 | 1:3000 | Rabbit |
| Cytokeratin 8 | Sigma-Aldrich #MABT329 | 1:1000 | Rat |
| CD3 | Abcam #ab11089 | 1:1000 | Rat |
| Ki67 | Thermo scientific #14-5698-82 | 1:1500 | Rat |
| Pan cytokeratin | Dako #3515 | 1:200 | Mouse |
| alpha Smooth muscle actin | Sigma-Aldrich #A5228 | 1:2000 | Mouse |

## References

Almagro, J., H.A. Messal, M. Zaw Thin, J. van Rheenen, and A. Behrens. 2021. Tissue clearing to examine tumour complexity in three dimensions. Nat Rev Cancer. 21:718–730. doi:10.1038/s41568-021-00382-w.

Anderson, T.J., D.J. Ferguson, and G.M. Raab. 1982. Cell turnover in the “resting” human breast: influence of parity, contraceptive pill, age and laterality. Br J Cancer. 46:376–382.

Arganda-Carreras, I., V. Kaynig, C. Rueden, K.W. Eliceiri, J. Schindelin, A. Cardona, and H. Sebastian Seung. 2017. Trainable Weka Segmentation: a machine learning tool for microscopy pixel classification. Bioinformatics. 33:2424–2426. doi:10.1093/bioinformatics/btx180.

Bankhead, P., M.B. Loughrey, J.A. Fernández, Y. Dombrowski, D.G. McArt, P.D. Dunne, S. McQuaid, R.T. Gray, L.J. Murray, H.G. Coleman, J.A. James, M. Salto-Tellez, and P.W. Hamilton. 2017. QuPath: Open source software for digital pathology image analysis. Sci Rep. 7:16878. doi:10.1038/s41598-017-17204-5.

Bannister, L., M. Berry, P. Collins, M. Dyson, J. Dussek, and M. Ferguson. 1995. Gray’s Anatomy: The anatomical basis of medicine and surgery. *In* Gray’s Anatomy. Churchill Livingstone, New York. 417–424.

Chen, J.-H., F. Liao, Y. Zhang, Y. Li, C.-J. Chang, C.-P. Chou, T.-L. Yang, and M.-Y. Su. 2017. 3D MRI for Quantitative Analysis of Quadrant Percent Breast Density: Correlation with Quadrant Location of Breast Cancer. Acad Radiol. 24:811–817. doi:10.1016/j.acra.2016.12.016.

Chen, Y., B. Pal, G.J. Lindeman, J.E. Visvader, and G.K. Smyth. 2022. R code and downstream analysis objects for the scRNA-seq atlas of normal and tumorigenic human breast tissue. Sci Data. 9:96. doi:10.1038/s41597-022-01236-2.

Cheng, N., N.A. Bhowmick, A. Chytil, A.E. Gorksa, K.A. Brown, R. Muraoka, C.L. Arteaga, E.G. Neilson, S.W. Hayward, and H.L. Moses. 2005. Loss of TGF-β type II receptor in fibroblasts promotes mammary carcinoma growth and invasion through upregulation of TGF-α-, MSP- and HGF-mediated signaling networks. Oncogene. 24:5053–5068. doi:10.1038/sj.onc.1208685.

Cheung, K.J., E. Gabrielson, Z. Werb, and A.J. Ewald. 2013. Collective Invasion in Breast Cancer Requires a Conserved Basal Epithelial Program. Cell. 155:1639–1651. doi:10.1016/j.cell.2013.11.029.

Dahl-Jensen, S.B., S. Yennek, L. Flasse, H.L. Larsen, D. Sever, G. Karremore, I. Novak, K. Sneppen, and A. Grapin-Botton. 2018. Deconstructing the principles of ductal network formation in the pancreas. PLOS Biology. 16:e2002842. doi:10.1371/journal.pbio.2002842.

van Dierendonck, J.H., R. Keijzer, C.J. van de Velde, and C.J. Cornelisse. 1989. Nuclear distribution of the Ki-67 antigen during the cell cycle: comparison with growth fraction in human breast cancer cells. Cancer Res. 49:2999–3006.

Dontu, G., and T.A. Ince. 2015. Of Mice and Women: A Comparative Tissue Biology Perspective of Breast Stem Cells and Differentiation. J Mammary Gland Biol Neoplasia. 20:51–62. doi:10.1007/s10911-015-9341-4.

Fendt, B.M., A. Hirschmann, M. Bruns, E. Camarillo-Retamosa, C. Ospelt, and A. Vogetseder. 2023. Protein atlas of fibroblast specific protein 1 (FSP1)/S100A4. Histol Histopathol. 38:1391– 1401. doi:10.14670/HH-18-621.

Figueroa, J.D., R.M. Pfeiffer, D.A. Patel, L. Linville, L.A. Brinton, G.L. Gierach, X.R. Yang, D. Papathomas, D. Visscher, C. Mies, A.C. Degnim, W.F. Anderson, S. Hewitt, Z.G. Khodr, S.E. Clare, A.M. Storniolo, and M.E. Sherman. 2014. Terminal Duct Lobular Unit Involution of the Normal Breast: Implications for Breast Cancer Etiology. J Natl Cancer Inst. 106:dju286. doi:10.1093/jnci/dju286.

Fujiwara, S., S. Deguchi, and T.M. Magin. 2020. Disease-associated keratin mutations reduce traction forces and compromise adhesion and collective migration. Journal of Cell Science. 133:jcs243956. doi:10.1242/jcs.243956.

Going, J.J., T.J. Anderson, S. Battersby, and C.C. MacIntyre. 1988. Proliferative and secretory activity in human breast during natural and artificial menstrual cycles. Am J Pathol. 130:193–204.

Goldhammer, N., J. Kim, R. Villadsen, L. Rønnov-Jessen, and O.W. Petersen. 2022. Myoepithelial progenitors as founder cells of hyperplastic human breast lesions upon PIK3CA transformation. Commun Biol. 5:1–13. doi:10.1038/s42003-022-03161-x.

Gouon-Evans, V., M.E. Rothenberg, and J.W. Pollard. 2000. Postnatal mammary gland development requires macrophages and eosinophils. Development. 127:2269–2282.

Gray, G.K., C.M.-C. Li, J.M. Rosenbluth, L.M. Selfors, N. Girnius, J.-R. Lin, R.C.J. Schackmann, W.L. Goh, K. Moore, H.K. Shapiro, S. Mei, K. D’Andrea, K.L. Nathanson, P.K. Sorger, S. Santagata, A. Regev, J.E. Garber, D.A. Dillon, and J.S. Brugge. 2022. A human breast atlas integrating single-cell proteomics and transcriptomics. Developmental Cell. 57:1400–1420.e7. doi:10.1016/j.devcel.2022.05.003.

Hannezo, E., C.L.G.J. Scheele, M. Moad, N. Drogo, R. Heer, R.V. Sampogna, J. van Rheenen, and B.D. Simons. 2017. A Unifying Theory of Branching Morphogenesis. Cell. 171:242–255.e27. doi:10.1016/j.cell.2017.08.026.

Hilton, H.N., C.L. Clarke, and J.D. Graham. 2018. Estrogen and progesterone signalling in the normal breast and its implications for cancer development. Molecular and Cellular Endocrinology. 466:2–14. doi:10.1016/j.mce.2017.08.011.

Howard, B.A., and B.A. Gusterson. 2000. Human Breast Development. J Mammary Gland Biol Neoplasia. 5:119–137. doi:10.1023/A:1026487120779.

Jindal, S., D. Gao, P. Bell, G. Albrektsen, S.M. Edgerton, C.B. Ambrosone, A.D. Thor, V.F. Borges, and P. Schedin. 2014. Postpartum breast involution reveals regression of secretory lobules mediated by tissue-remodeling. Breast Cancer Research. 16:R31. doi:10.1186/bcr3633.

Kohler, K.T., N. Goldhammer, S. Demharter, U. Pfisterer, K. Khodosevich, L. Rønnov-Jessen, O.W. Petersen, R. Villadsen, and J. Kim. 2022. Ductal keratin 15+ luminal progenitors in normal breast exhibit a basal-like breast cancer transcriptomic signature. npj Breast Cancer. 8:1–12. doi:10.1038/s41523-022-00444-8.

Koledova, Z., X. Zhang, C. Streuli, R.B. Clarke, O.D. Klein, Z. Werb, and P. Lu. 2016. SPRY1 regulates mammary epithelial morphogenesis by modulating EGFR-dependent stromal paracrine signaling and ECM remodeling. Proc Natl Acad Sci U S A. 113:E5731–E5740. doi:10.1073/pnas.1611532113.

Kumar, T., K. Nee, R. Wei, S. He, Q.H. Nguyen, S. Bai, K. Blake, M. Pein, Y. Gong, E. Sei, M. Hu, A.K. Casasent, A. Thennavan, J. Li, T. Tran, K. Chen, B. Nilges, N. Kashikar, O. Braubach, B. Ben Cheikh, N. Nikulina, H. Chen, M. Teshome, B. Menegaz, H. Javaid, C. Nagi, J. Montalvan, T. Lev, S. Mallya, D.F. Tifrea, R. Edwards, E. Lin, R. Parajuli, S. Hanson, S. Winocour, A. Thompson, B. Lim, D.A. Lawson, K. Kessenbrock, and N. Navin. 2023. A spatially resolved single-cell genomic atlas of the adult human breast. Nature. 620:181–191. doi:10.1038/s41586-023-06252-9.

Laidlaw, I.J., R.B. Clarke, A. Howell, A.W. Owen, C.S. Potten, and E. Anderson. 1995. The proliferation of normal human breast tissue implanted into athymic nude mice is stimulated by estrogen but not progesterone. Endocrinology. 136:164–171. doi:10.1210/endo.136.1.7828527.

Le Hir, M., I. Hegyi, D. Cueni-Loffing, J. Loffing, and B. Kaissling. 2005. Characterization of renal interstitial fibroblast-specific protein 1/S100A4-positive cells in healthy and inflamed rodent kidneys. Histochem Cell Biol. 123:335–346. doi:10.1007/s00418-005-0788-z.

Lefevre, J.G., K.M. Short, T.O. Lamberton, O. Michos, D. Graf, I.M. Smyth, and N.A. Hamilton. 2017. Branching morphogenesis in the developing kidney is governed by rules that pattern the ureteric tree. Development. 144:4377–4385. doi:10.1242/dev.153874.

Longacre, T.A., and S.A. Bartow. 1986. A correlative morphologic study of human breast and endometrium in the menstrual cycle. Am J Surg Pathol. 10:382–393. doi:10.1097/00000478-198606000-00003.

Macias, H., and L. Hinck. 2012. Mammary gland development. WIREs Developmental Biology. 1:533–557. doi:10.1002/wdev.35.

Messal, H.A., J. Almagro, M. Zaw Thin, A. Tedeschi, A. Ciccarelli, L. Blackie, K.I. Anderson, I. Miguel-Aliaga, J. van Rheenen, and A. Behrens. 2021. Antigen retrieval and clearing for whole-organ immunofluorescence by FLASH. Nature Protocols. 16:239–262. doi:10.1038/s41596-020-00414-z.

Metzger, R.J., O.D. Klein, G.R. Martin, and M.A. Krasnow. 2008. The branching programme of mouse lung development. Nature. 453:745–750. doi:10.1038/nature07005.

Morsing, M., M.C. Klitgaard, A. Jafari, R. Villadsen, M. Kassem, O.W. Petersen, and L. Rønnov-Jessen. 2016. Evidence of two distinct functionally specialized fibroblast lineages in breast stroma. Breast Cancer Res. 18:108. doi:10.1186/s13058-016-0769-2.

Nelson, C.M., M.M. VanDuijn, J.L. Inman, D.A. Fletcher, and M.J. Bissell. 2006. Tissue Geometry Determines Sites of Mammary Branching Morphogenesis in Organotypic Cultures. Science. 314:298–300. doi:10.1126/science.1131000.

Nerger, B.A., J.M. Jaslove, H.E. Elashal, S. Mao, A. Košmrlj, A.J. Link, and C.M. Nelson. 2021. Local accumulation of extracellular matrix regulates global morphogenetic patterning in the developing mammary gland. Curr Biol. 31:1903–1917.e6. doi:10.1016/j.cub.2021.02.015

Nguyen, Q.H., N. Pervolarakis, K. Blake, D. Ma, R.T. Davis, N. James, A.T. Phung, E. Willey, R. Kumar, E. Jabart, I. Driver, J. Rock, A. Goga, S.A. Khan, D.A. Lawson, Z. Werb, and K. Kessenbrock. 2018. Profiling human breast epithelial cells using single cell RNA sequencing identifies cell diversity. Nature Communications. 9:1–12. doi:10.1038/s41467-018-04334-1.

Ogony, J., T. de Bel, D.C. Radisky, J. Kachergus, E.A. Thompson, A.C. Degnim, K.J. Ruddy, T. Hilton, M. Stallings-Mann, C. Vachon, T.L. Hoskin, M.G. Heckman, R.A. Vierkant, L.J. White, R.M. Moore, J. Carter, M. Jensen, L. Pacheco-Spann, J.E. Henry, A.M. Storniolo, S.J. Winham, J. van der Laak, and M.E. Sherman. 2022. Towards defining morphologic parameters of normal parous and nulliparous breast tissues by artificial intelligence. Breast Cancer Res. 24:45. doi:10.1186/s13058-022-01541-z.

Osin, P.P., R. Anbazhagan, J. Bartkova, B. Nathan, and B.A. Gusterson. 1998. Breast development gives insights into breast disease. Histopathology. 33:275–283. doi:10.1046/j.1365-2559.1998.00479.x.

Österreicher, C.H., M. Penz-Österreicher, S.I. Grivennikov, M. Guma, E.K. Koltsova, C. Datz, R. Sasik, G. Hardiman, M. Karin, and D.A. Brenner. 2011. Fibroblast-specific protein 1 identifies an inflammatory subpopulation of macrophages in the liver. Proc Natl Acad Sci U S A. 108:308–313. doi:10.1073/pnas.1017547108.

Pal, B., Y. Chen, F. Vaillant, B.D. Capaldo, R. Joyce, X. Song, V.L. Bryant, J.S. Penington, L. Di Stefano, N. Tubau Ribera, S. Wilcox, G.B. Mann, kConFab, A.T. Papenfuss, G.J. Lindeman, G.K. Smyth, and J.E. Visvader. 2021. A single-cell RNA expression atlas of normal, preneoplastic and tumorigenic states in the human breast. EMBO J. 40:e107333. doi:10.15252/embj.2020107333.

Perinatal statistics - parturients, delivers and newborns - THL. 2024. Finnish Institute for Health and Welfare (THL), Finland.

Peuhu, E., R. Kaukonen, M. Lerche, M. Saari, C. Guzmán, P. Rantakari, N. De Franceschi, A. Wärri, M. Georgiadou, G. Jacquemet, E. Mattila, R. Virtakoivu, Y. Liu, Y. Attieh, K.A. Silva, T. Betz, J.P. Sundberg, M. Salmi, M.-A. Deugnier, K.W. Eliceiri, and J. Ivaska. 2017. SHARPIN regulates collagen architecture and ductal outgrowth in the developing mouse mammary gland. EMBO J. 36:165–182. doi:10.15252/embj.201694387.

Peurla, M. 2023. 3D branching simulation v. 3.21. 1. doi:10.17632/y8d52ygdxv.1.

Pickup, M.W., L.D. Hover, E.R. Polikowsky, A. Chytil, A.E. Gorska, S.V. Novitskiy, H.L. Moses, and P. Owens. 2015. BMPR2 loss in fibroblasts promotes mammary carcinoma metastasis via increased inflammation. Molecular Oncology. 9:179–191. doi:10.1016/j.molonc.2014.08.004.

Potten, C.S., R.J. Watson, G.T. Williams, S. Tickle, S.A. Roberts, M. Harris, and A. Howell. 1988. The effect of age and menstrual cycle upon proliferative activity of the normal human breast. Br J Cancer. 58:163–170. doi:10.1038/bjc.1988.185.

Prabhu, S.D., H.S. Rai, R. Nayak, R. Naik, and S. Jayasheelan. Study of the Immunohistochemical Expression of p63 in Benign Lesions and Carcinoma of the Breast at a Tertiary Hospital in South India. Cureus. 15:e48557. doi:10.7759/cureus.48557.

Ramakrishnan, R., S.A. Khan, and S. Badve. 2002. Morphological Changes in Breast Tissue with Menstrual Cycle. Mod Pathol. 15:1348–1356. doi:10.1097/01.MP.0000039566.20817.46.

Ramms, L., G. Fabris, R. Windoffer, N. Schwarz, R. Springer, C. Zhou, J. Lazar, S. Stiefel, N. Hersch, U. Schnakenberg, T.M. Magin, R.E. Leube, R. Merkel, and B. Hoffmann. 2013. Keratins as the main component for the mechanical integrity of keratinocytes. Proc Natl Acad Sci U S A. 110:18513–18518. doi:10.1073/pnas.1313491110.

Rios, A.C., B.D. Capaldo, F. Vaillant, B. Pal, R. van Ineveld, C.A. Dawson, Y. Chen, E. Nolan, N.Y. Fu, 3DTCLSM Group, F.C. Jackling, S. Devi, D. Clouston, L. Whitehead, G.K. Smyth, S.N. Mueller, G.J. Lindeman, and J.E. Visvader. 2019. Intraclonal Plasticity in Mammary Tumors Revealed through Large-Scale Single-Cell Resolution 3D Imaging. Cancer Cell. 35:618–632.e6. doi:10.1016/j.ccell.2019.02.010.

Roos, F.J.M., G.S. van Tienderen, H. Wu, I. Bordeu, D. Vinke, L.M. Albarinos, K. Monfils, S. Niesten, R. Smits, J. Willemse, O. Rosmark, G. Westergren-Thorsson, D.J. Kunz, M. de Wit, P.J. French, L. Vallier, J.N.M. IJzermans, R. Bartfai, H. Marks, B.D. Simons, M.E. van Royen, M.M.A. Verstegen, and L.J.W. van der Laan. 2022. Human branching cholangiocyte organoids recapitulate functional bile duct formation. Cell Stem Cell. 29:776–794.e13. doi:10.1016/j.stem.2022.04.011.

Rosebrock, A., J.J. Caban, J. Figueroa, G. Gierach, L. Linville, S. Hewitt, and M. Sherman. 2013. Quantitative Analysis of TDLUs using Adaptive Morphological Shape Techniques. Proc SPIE. 8676:86760N. doi:10.1117/12.2006619.

Russo, J., X. Ao, C. Grill, and I.H. Russo. 1999. Pattern of distribution of cells positive for estrogen receptor α and progesterone receptor in relation to proliferating cells in the mammary gland. Breast Cancer Res Treat. 53:217–227. doi:10.1023/A:1006186719322.

Russo, J., R. Rivera, and I.H. Russo. 1992. Influence of age and parity on the development of the human breast. Breast Cancer Res Treat. 23:211–218. doi:10.1007/BF01833517.

Russo, J., A.L. Romero, and I.H. Russo. 1994. Architectural pattern of the normal and cancerous breast under the influence of parity. Cancer Epidemiol Biomarkers Prev. 3:219–224.

Russo, J., and I.H. Russo. 1997. Differentiation and breast cancer. Medicina (B Aires). 57 Suppl 2:81– 91.

Ruusuvuori, P., M. Valkonen, K. Kartasalo, M. Valkonen, T. Visakorpi, M. Nykter, and L. Latonen. 2022. Spatial analysis of histology in 3D: quantification and visualization of organ and tumor level tissue environment. Heliyon. 8:e08762. doi:10.1016/j.heliyon.2022.e08762.

Santandrea, G., C. Bellarosa, D. Gibertoni, M.C. Cucchi, A.M. Sanchez, G. Franceschini, R. Masetti, and M.P. Foschini. 2021. Hormone Receptor Expression Variations in Normal Breast Tissue: Preliminary Results of a Prospective Observational Study. J Pers Med. 11:387. doi:10.3390/jpm11050387.

Schindelin, J., I. Arganda-Carreras, E. Frise, V. Kaynig, M. Longair, T. Pietzsch, S. Preibisch, C. Rueden S. Saalfeld, B. Schmid, J.-Y. Tinevez, D.J. White, V. Hartenstein, K. Eliceiri, P. Tomancak, and A. Cardona. 2012. Fiji: an open-source platform for biological-image analysis. Nat Methods. 9:676–682. doi:10.1038/nmeth.2019.

Shah, A., G. Haider, N. Abro, S. Bhutto, T.I. Baqai, S. Akhtar, and K. Abbas. 2022. Correlation Between Age and Hormone Receptor Status in Women With Breast Cancer. Cureus. 14:e21652. doi:10.7759/cureus.21652.

Shoker, B.S., C. Jarvis, D.R. Sibson, C. Walker, and J.P. Sloane. 1999. Oestrogen receptor expression in the normal and pre-cancerous breast. The Journal of Pathology. 188:237–244. doi:10.1002/(SICI)1096-9896(199907)188:3<237::AID-PATH343>3.0.CO;2-8.

Short, K., M. Hodson, and I. Smyth. 2013. Spatial mapping and quantification of developmental branching morphogenesis. Development. 140:471–478. doi:10.1242/dev.088500.

Silberstein, G.B. 2001. Postnatal mammary gland morphogenesis. Microscopy Research and Technique. 52:155–162. doi:10.1002/1097-0029(20010115)52:2<155::AID-JEMT1001>3.0.CO;2-P.

Silberstein, G.B., and C.W. Daniel. 1984. Glycosaminoglycans in the basal lamina and extracellular matrix of serially aged mouse mammary ducts. Mech Ageing Dev. 24:151–162. doi:10.1016/0047-6374(84)90067-8.

Silberstein, G.B., and C.W. Daniel. 1987. Reversible Inhibition of Mammary Gland Growth by Transforming Growth Factor-β. Science. 237:291–293. doi:10.1126/science.3474783.

Söderqvist, G., E. Isaksson, B. von Schoultz, K. Carlström, E. Tani, and L. Skoog. 1997. Proliferation of breast epithelial cells in healthy women during the menstrual cycle. American Journal of Obstetrics & Gynecology. 176:123–128. doi:10.1016/S0002-9378(97)80024-5.

Sumbal, J., S. Fre, and Z.S. Koledova. 2024. Fibroblast-induced mammary epithelial branching depends on fibroblast contractility. PLOS Biology. 22:e3002093. doi:10.1371/journal.pbio.3002093.

Sumbal, J., and Z. Koledova. 2019. FGF signaling in mammary gland fibroblasts regulates multiple fibroblast functions and mammary epithelial morphogenesis. Development. 146:dev185306. doi:10.1242/dev.185306.

Susaki, E.A., K. Tainaka, D. Perrin, F. Kishino, T. Tawara, T.M. Watanabe, C. Yokoyama, H. Onoe, M. Eguchi, S. Yamaguchi, T. Abe, H. Kiyonari, Y. Shimizu, A. Miyawaki, H. Yokota, and H.R. Ueda. 2014. Whole-Brain Imaging with Single-Cell Resolution Using Chemical Cocktails and Computational Analysis. Cell. 157:726–739. doi:10.1016/j.cell.2014.03.042.

Sznurkowska, M.K., E. Hannezo, R. Azzarelli, S. Rulands, S. Nestorowa, C.J. Hindley, J. Nichols, B. Göttgens, M. Huch, A. Philpott, and B.D. Simons. 2018. Defining Lineage Potential and Fate Behavior of Precursors during Pancreas Development. Dev Cell. 46:360–375.e5. doi:10.1016/j.devcel.2018.06.028.

Tabár, L., P.B. Dean, F.L. Tucker, A.M.-F. Yen, J.C.-Y. Fann, A.T.-Y. Lin, R.A. Smith, S.W. Duffy, and T.H.-H. Chen. 2022. Breast cancers originating from the terminal ductal lobular units: In situ and invasive acinar adenocarcinoma of the breast, AAB. European Journal of Radiology. 152:110323. doi:10.1016/j.ejrad.2022.110323.

Taylor-Papadimitriou, J., M. Stampfer, J. Bartek, A. Lewis, M. Boshell, E.B. Lane, and I.M. Leigh. 1989. Keratin expression in human mammary epithelial cells cultured from normal and malignant tissue: relation to in vivo phenotypes and influence of medium. Journal of Cell Science. 94:403–413. doi:10.1242/jcs.94.3.403.

Trimboli, A.J., C.Z. Cantemir-Stone, F. Li, J.A. Wallace, A. Merchant, N. Creasap, J.C. Thompson, E. Caserta, H. Wang, J.-L. Chong, S. Naidu, G. Wei, S.M. Sharma, J.A. Stephens, S.A. Fernandez, M.N. Gurcan, M.B. Weinstein, S.H. Barsky, L. Yee, T.J. Rosol, P.C. Stromberg, M.L. Robinson, F. Pepin, M. Hallett, M. Park, M.C. Ostrowski, and G. Leone. 2009. Pten in Stromal Fibroblasts Suppresses Mammary Epithelial Tumors. Nature. 461:1084–1091. doi:10.1038/nature08486.

Uçar, M.C., D. Kamenev, K. Sunadome, D. Fachet, F. Lallemend, I. Adameyko, S. Hadjab, and E. Hannezo. 2021. Theory of branching morphogenesis by local interactions and global guidance. Nat Commun. 12:6830. doi:10.1038/s41467-021-27135-5.

Villadsen, R., A.J. Fridriksdottir, L. Rønnov-Jessen, T. Gudjonsson, F. Rank, M.A. LaBarge, M.J. Bissell, and O.W. Petersen. 2007. Evidence for a stem cell hierarchy in the adult human breast. Journal of Cell Biology. 177:87–101. doi:10.1083/jcb.200611114.

Vince, J. 2021. Vector Analysis for Computer Graphics. Springer, London.

Wellings, S.R., H.M. Jensen, and R.G. Marcum. 1975. An Atlas of Subgross Pathology of the Human Breast With Special Reference to Possible Precancerous Lesions2. JNCI: Journal of the National Cancer Institute. 55:231–273. doi:10.1093/jnci/55.2.231.

Williams, G., E. Anderson, A. Howell, R. Watson, J. Covne, S.A. Roberts, and C.S. Potten. 1991. Oral contraceptive (OCP) use increases proliferation and decreases oestrogen receptor content of epithelial cells in the normal human breast. International Journal of Cancer. 48:206–210. doi:10.1002/ijc.2910480209.

Woodford, O. 2024. vol3d v2 - File Exchange - MATLAB Central.

Zhao, C., S. Cai, K. Shin, A. Lim, T. Kalisky, W.-J. Lu, M.F. Clarke, and P.A. Beachy. 2017. Stromal Gli2 activity coordinates a niche signaling program for mammary epithelial stem cells. Science. 356:eaal3485. doi:10.1126/science.aal3485.

